# An experimental model system to investigate microscale mechanisms behind the soil priming effect

**DOI:** 10.64898/2026.07.14.738410

**Authors:** Moritz Mohrlok, Christina Kaiser

## Abstract

The soil priming effect plays an important role in the global carbon cycle. Although well-studied, the mechanisms behind it remain elusive. So far, studies measured the phenomenon at the bulk soil scale, neglecting that it arises from spatially explicit processes that take place at the microscale. Here, we present a novel approach using a microfluidic device to directly assess the response of soil microbes living on a patch of complex substrate to a pulse of easily available substrate. Using time-resolved fluorescence microscopy, we tracked motility, position, shape characteristics and attached biomass of green fluorescent protein expressing *Bacillus subtilis* cells living on a transparent carboxymethylcellulose substrate patch exposed to a pulse of growth medium with differing concentrations. Assessing CMC decomposition via Congo-Red staining after 42 days of incubations with constant observation, we observed increased decomposition upon addition of enough labile substrate, resembling a priming effect. The pulse triggered a transient increase in bacterial motility, indicating the formation of exploring and growing subpopulations respectively. We observed a concentration-dependent growth response, resulting in different behaviors. High concentrations led to high biomass and decomposition of CMC, however growth was quickly limited, possibly by depletion of necessary nutrients and waste accumulation. Intermediate concentration, however, resulted in a more sustained attached biomass, that showed evidence of spatial self-organization, leading to similar decomposition. We present a novel experimental model system to study the behavior of soil microbes decomposing complex substrate and provide a unique view into the response of such a population to a labile substrate pulse.

## Introduction

Soils store vast amount of carbon (Lal & Stewart, 2019), whose fate relies on the action of microorganisms inhabiting this patchy and heterogenous habitat (Lehmann et al., 2020). They take up and metabolize easily available, low molecular weight compounds but are also able to degrade more complex soil organic matter (SOM) through the production of extracellular enzymes. Organic matter can get stabilized or respired depending on the activity of these microbes, playing an important role in the terrestrial carbon cycle (Sharma et al., 2025).

The addition of easily available OM to soil often triggers rapid growth of microbial biomass and a peak of respiration. This respired carbon does not only stem from the added OM, but also from stable, older SOM. The increase in mineralization of SOM through labile OM addition has been termed the positive priming effect (Bernard et al., 2022; Kuzyakov, 2010). As this additional release of stabilized C could have strong implications for the global carbon cycle (Zhang et al., 2025), the priming effect has been the subject to a plethora of studies in the last two decades. Different mechanisms have been proposed that could each play a role depending on the nutrient status of soil, stoichiometry of fresh and stable substrate and microbial growth strategies (Bernard 2022). For example, fresh OM input can increase SOM decomposition through the “microbial activation” mechanism by generally increasing microbial biomass and enzymatic activity (Bernard et al., 2022; Chen et al., 2014; Liu et al., 2020), and by alleviating the energy limitation of soil microbes (Bernard et al., 2022; Blagodatskaya & Kuzyakov, 2008; Kuzyakov et al., 2000; Lehmann et al., 2020; Yu et al., 2018). Microbes first utilize the fresh labile substrate, and then the increased and more active biomass turns to older SOM (Liu et al., 2020). An important factor is the amount of added OM. Evidence exists that a certain minimum addition is necessary to trigger priming (Liu et al., 2017). On the other hand, a review by Blagodatskaya & Kuzyakov (2008) found a negative concentration-dependence of priming when added substrate exceeded 50% of microbial biomass C, suggesting that microbes preferentially using the vast amount of labile substrate instead of older SOM. Direct experiments by Di Lonardo et al. (2019) and Liu et al. (2017) however showed that this pattern is not universal, and priming can increase linearly with substrate concentration under certain conditions.

These examples highlight the complexity of the topic, resulting in often contrasting outcomes in different soils (Zhang et al., 2025). While a lot of progress has been made in understanding the priming effect, most studies are bulk soil approaches, that measure at a spatially aggregated level. The priming effect, however, emerges from spatially explicit processes that happen at the scale of soil pores, substrate particles, individual microbes and colonies, the microscale. For example, a possible mechanism to our knowledge only described in marine systems so far (Ebrahimi et al., 2019) but also similarly observed in a soil-focused individual-based model (Kaiser et al., 2017), is density-dependency of complex substrate degradation. The degradation products of extracellular enzymes can diffuse away from a SOM-degrading microbial colony before being taken up, meaning the return of investment into costly enzymes is reduced. An increase of microbial density upon the addition of enough labile substrate could alleviate this limitation as it increases the likelihood that diffusing products can be taken up by a community member. This increases decomposition efficiency and could allow more sustained degradation.

A direct understanding of such foundational mechanisms might be the key to understanding the priming effect. There is currently a lack of studies on such microscale processes in soil, as direct observations of how microbes react to for example labile substrate pulses are limited by the opacity of soil and the small size of microorganisms (Zhu et al., 2022). A possible solution are microfluidic devices, which have been increasingly used as experimental model systems in soil microbial ecology (Rusconi et al., 2014; Zhu et al., 2022). These devices allow the direct observation of microbes in a controlled, reduced-complexity environment with similar properties as soil (Aleklett et al., 2018; Arellano-Caicedo et al., 2021; Zhu et al., 2022). Such experimental model systems have not yet been utilized to study the priming effect.

While microfluidic systems exist that allow the observation of bacteria during the extracellular enzyme-requiring decomposition of polymers like xylose (D’Souza et al., 2021) and could be adapted to study priming, these systems lack the spatial aspect of SOM decomposition as the substrate is dissolved and itself subject to diffusion. Here, we present a prototypical experimental model system that allows the direct study of fluorescently labelled microorganisms growing on a solid, transparent patch of complex substrate. Specifically, we incubated millimeter-sized chambers containing a transparent carboxymethylcellulose-agarose gel as complex carbon source with a green-fluorescent protein (GFP)-expressing strain of the soil bacterium *Bacillus subtilis*. We simulated a priming event by filling the chamber with different concentrations of growth medium (LB), containing a mix of low molecular weight organic compounds that the strain grows well on, and tracked bacterial biomass, position and motility using fluorescence microscopy and a custom image analysis pipeline at multiple timepoints over the course of 42 days. CMC decomposition was finally assessed by Congo Red staining of the remaining substrate patches and subsequent analysis of their fluorescence intensity. Our results allow us to visualize the direct response of a bacterial community to a labile substrate pulse, and the effect that this has on complex substrate degradation.

The main research question of this study is whether we can simulate the priming effect in the developed experimental model system. We expect the bacteria to decompose more complex substrate when exposed to a higher labile substrate pulse. Specifically, we hypothesize that the substrate pulse leads to increased biomass, scaling with substrate concentration, which leads to higher decomposition. The secondary research question is whether we can identify microscale mechanisms that play a role behind priming in this system. We hypothesize to observe bacterial colonies that cross a density threshold upon sufficient nutrient addition, allowing sustained growth on CMC once labile substrate is depleted.

## Material & Methods

Figure 1 provides an overview of the experimental workflow, from chip fabrication and inoculation, time-series imaging and the assessment of CMC decomposition to image analysis. Each step is described in the following, with a more detailed description provided in the supplement (S14).

**Figure 1:**
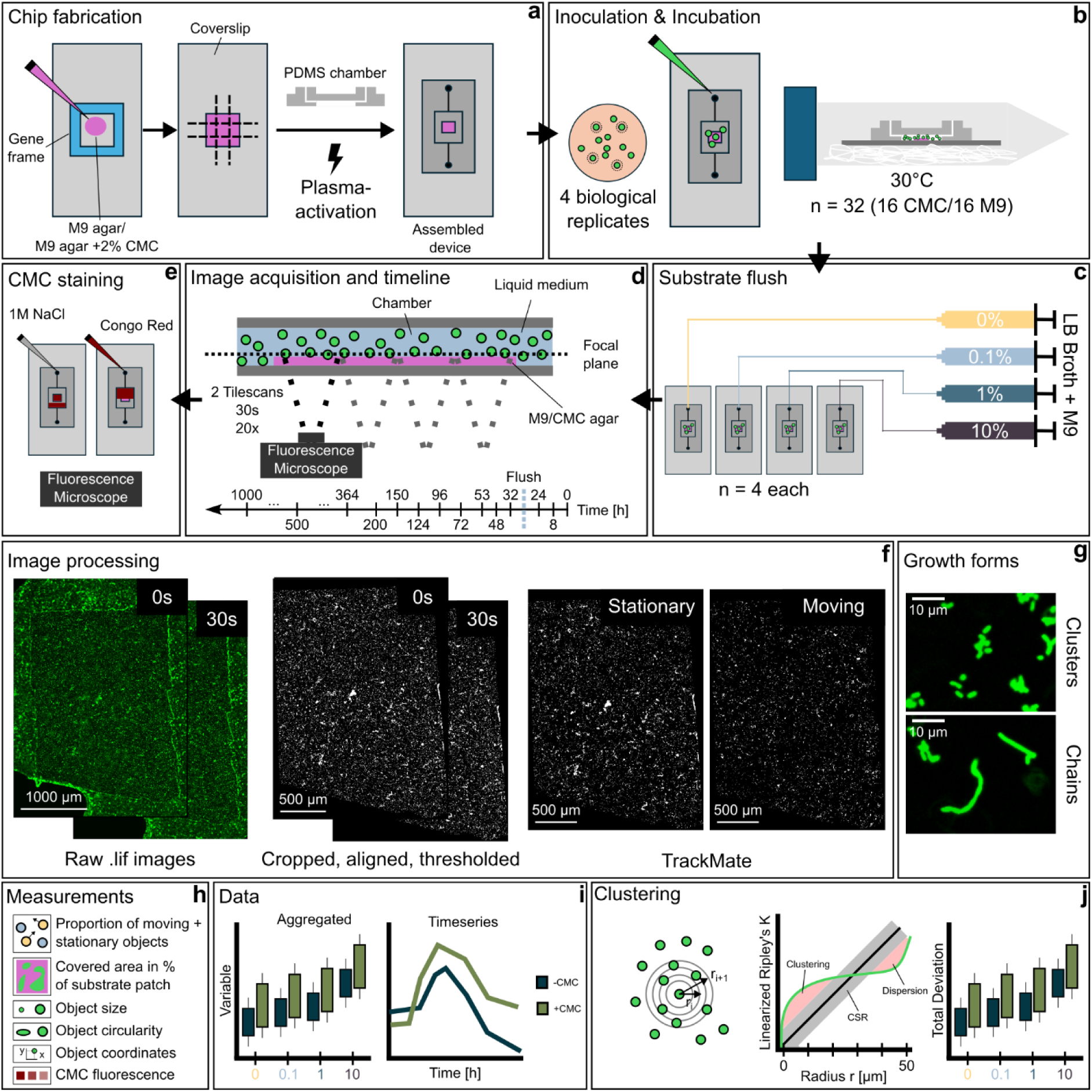
Overview of the methodology developed and employed in this study. Substrate patches were fabricated by pipetting warm M9 or M9 + 2% carboxymethylcellulose-agar (CMC) into gene frames attached to glass coverslips and then removing excess substrate. Microfluidic chambers were constructed by attaching a PDMS chamber on top of the substrate patch using a plasma oven (**a**). GFP-expressing B. Subtilis was pre-cultured on LB agar plates supplemented with 2% CMC. For each biological replicate (4 in total), one colony was suspended in M9 minimal media supplemented with 0.1% LB, diluted to the same OD_600_ (0.001) and then carefully pipetted in the chip. Inoculated chips (4 LB x 2 CMC treatments per replicate = 32 chips in total) were sealed and incubated at 30 °C horizontally in falcon tubes containing a wet paper towel (**b**). After an acclimatization period, chips were attached to syringe pumps filled with different concentrations of LB growth medium diluted in M9 minimal medium (**c**) and incubated again. Imaging was done during the incubation at multiple timepoints (**d**) Two tilescans 30 seconds apart were taken at every timepoint using an inverted fluorescence microscope. After the experiment, chips were etOH-fixed, stained with Congo Red and washed with NaCl. Fluorescence intensity (575/590 nm) was then recorded to assess CMC decomposition (lower fluorescence = less CMC left over = higher decomposition) (**e**). Raw images were cropped to the substrate area, aligned and thresholded to differentiate background and cells in FIJI/ImageJ. The image shown is representative of a highly crowded image taken shortly after the substrate pulse (Chip 16, +CMC/ 10% LB). Moving were separated from stationary objects using the TrackMate plugin across the two images taken at every timepoint (**f**). Cells presented as clusters of individual cells or formed chains during growth (**g**). Different measurements were derived from processed images (**h**) and analyzed in different ways (**i**). Positions of cells in two-dimensional space were used to calculate the linearized form of Ripley’s K function with radius r = 50 µm, comparing it to complete spatial randomness (CSR) giving an estimate of spatial cell clustering (**j**).

### Bacteria and culture conditions

GFP-expressing *Bacillus subtilis* strain 4819 was pre-grown at 28 °C on LB agar plates supplemented with 2% w/v CMC (Sigma-Aldrich). M9 minimal media was prepared according to Souza et al. (2009) and supplemented with 20 mg * l^-1^ tryptophane (Sigma-Aldrich). Without it, the strain was unable to grow in M9 supplemented with other carbon sources like glucose in pre-experiments. Tryptophane alone however did not trigger a strong growth response (see the-CMC/0% LB treatment in the results). For the substrate patches, CMC (2% w/v) was dissolved in M9, and agarose (1.5% w/v, Roth) was added before autoclaving. A M9 + agarose gel without CMC was similarly prepared for the -CMC treatment. LB medium was prepared according to standard protocol and diluted in M9 to 0.1, 1 and 10% (v/v) for the priming treatments. This medium was chosen because it triggered a consistent and strong growth response of the strain, providing all essential nutrients and energy it needs. All culturing work, and handling of inoculated unsealed chips was performed in a laminar flow hood using aseptic technique. Inoculated, sealed chips were incubated horizontally in closed falcon tubes with wet tissue paper to prevent drying out at 30 °C when not being observed under the microscope.

### Chip fabrication

The microfluidic chips consisted of a glass coverslip (24 x 60 mm, Marienfeld), a patch of M9-agarose substrate (with or without CMC) and a chamber made from polydimethylsiloxane (PDMS, Sylgard). These were fabricated from the same microfluidic master wafer as used by Sharma et al. (2020). Ten µl of dissolved, warm agarose medium was pipetted onto gene frames (Thermo Scientifc, 1×1 cm). After scoring the solidified gel with a scalpel in a checkerboard pattern and removing the gene frame and excess substrate, a rectangular substrate patch of about 2 x 4 mm remained on the coverslip. The PDMS chamber with in- and outlets was then manually aligned with the patch and bonded to the glass using a plasma oven (Zepto MHz, Diener electronic). Finished chips were sterilized using 70% etOH and flushed with M9 for 24 hours using syringe pumps. In-and outlets of liquid-filled chips were sealed with small strips of autoclaved aluminum foil and transparent sticky tape and stored as described above.

### Experimental setup and incubation

Four different colonies of *B. subtilis* were harvested and each diluted in M9 containing 0.1% LB to an OD_600_ of 0.001. Treating these colonies as biological replicates, 4 chips with and without CMC were inoculated per replicate, resulting in 16 chips per CMC treatment. After a 48-hour acclimatization period, a labile substrate pulse was simulated by replacing the chamber volumes of each biological replicate with either M9 (= 0% LB), 0.1, 1 or 10% LB using a 12-channel syringe pump (SPLab12, Baoding Shenchen) set to a minimal flow rate (0.7 µl * min^-1^) to minimize bacterial displacement. After priming, chips were sealed again and incubated at 30 °C.

### Image acquisition

Chips were taken out of their incubation tubes and imaged at multiple timepoints before and after the priming pulse. Images were acquired using a wide-field inverted fluorescence microscope (Thunder Imager, Leica Microsystems) using a 20x air objective (HC PL Fluotar 20x/0.55, Leica Microsystems). A tilescan of the whole agarose patch was acquired at each timepoint, focusing on the first visible layer of bacteria, which were those sitting directly on the substrate. Baseline imaging settings were optimized for speed and image quality, using 200 ms of exposure time with 100% light intensity (475/535 nm excitation/emission). Exposure time and light intensity were adjusted to avoid overexposure when cell density was high. Two images with identical settings were taken with a 30 second interval between acquisition at each timepoint. After imaging, chips were returned to their tubes and the incubator.

### Image processing

All image processing was done in Fiji (2.16.0/1.54p) with custom, semi-automated scripts to ensure the same treatment of each image. Images were converted to 8 bit to reduce file size. Images were aligned to the timepoint 1 image of each chip using the “line ROI alignment” tool. Each image was then cropped to the substrate patch area. After applying a Gaussian blur filter (σ = 2), images were binarized by grouping images by their set of image acquisition parameters (light intensity + exposure time) and applying the Otsu thresholding algorithm to find an automatic threshold for each image, subsequently applying the average threshold to all images of each group. Artifacts (objects that did not change over time, likely auto-fluorescent dust on the outside of the chambers) were manually deleted from some images. Making use of the two images at each timepoint, we were able to separate each image into moving and stationary objects using the TrackMate plugin (7.14.10, Ershov et al., 2022). Settings are described in more detail in the supplement. We then extracted the moving and stationary object count, covered area and average circularity for each chip and timepoint. In addition, a list of all objects of each image with size, circularity and coordinates was exported.

### CMC staining

Congo Red (CR) was used to stain the agarose patches after the experiment, as this dye increases its fluorescence upon binding to CMC (Zemanek et al., 2017) and can thus be used to assess CMC decomposition (Romano et al., 2013). Chips were fixed with 96% etOH, stained with CR followed by 1M NaCl and washed multiple times with MQ. Stained gels were then imaged in the same way as before, with constant imaging settings (575/590 nm excitation/emission, 200 ms exposure time with 100% light intensity). Images were cropped to the substrate patch area, down-sampled by 50% to reduce file size, and fluorescence intensity was extracted for each pixel.

### Calculations and statistics

All calculations and statistics were done in R (4.4.3, R Core Team (2025) using RStudio (2025.09.1, Posit Software). Moving and stationary object counts were normalized to the observed patch area. Similarly, total area covered by stationary bacteria was expressed in percent of the patch area in each image. The proportion of moving and stationary objects was calculated as percentage of the total objects. The log-response ratio (LRR) of normalized moving and stationary object counts as well as covered area was calculated relative to the average of -CMC/0% LB chips for each timepoint, adding a small constant value to each datapoint due to zeros in the data. Average fluorescence intensity of the Congo Red-stained images was calculated from all pixels of each down-sampled image. This value was further normalized by the average stationary biomass-covered area over the entire time-series. Standard statistical tests are mentioned when applicable and further information on them is provided in the supplement.

The effect of LB concentration and averaged measurements over the entire time-series on the final CR fluorescence values was assessed using linear modelling (S11). By stepwise model reduction based on the Akaike Information Criterion (AIC), we obtained the best predictors for CMC decomposition.

Time-series were analyzed using generalized additive models with R package “mgcv” (1.9-1, Wood, 2017). CMC and LB concentration were included as linear predictors; time was added as a smooth term with cubic regression splines (S2). To analyze the change of LRRs over time, we fitted a model for each LB level, using only CMC as predictor. Random splines were added for each chip to control for inherent differences. Partial and marginal effects were obtained to quantify the effects of the non-smooth model terms while keeping all other terms constant or considering the whole model respectively. To compare treatments, the timespans where curves differed significantly from each other were computed via difference smooths.

Object coordinates were used to assess spatial organization of bacteria (S9) using R package “spatstat” (Baddeley et al., 2015). We calculated the linearized form of Ripley’s K (Ripley, 1988) for each point in two-dimensional space with a radius r of 50 µm, which we deemed a reasonable distance given the average interaction distance of soil microbes being about 20 µm (Raynaud & Nunan, 2014). This approach compares the observed number of objects in a specified radius around an object to the average expected number under complete spatial randomness (CSR) and provides evidence for significant clustering or dispersion (Raynaud & Nunan, 2014). We quantified this deviance as the difference between the observed and expected curve across r (see Fig. 1j) and assessed significant deviations form CSR with Diggle-Cressie-Loosmore-Ford tests (Baddeley et al., 2014). Bacterial clusters formed large, coherent objects in our data that would be represented by only one coordinate as our microscopy approach did not allow discerning individual cells in these clusters, affecting this analysis. To circumvent this, we approximated the bacterial distribution within large objects (larger than one standard deviation from the average size) using data of higher resolution images of single colonies taking during the experiment (Fig. S8).

## Results

We investigated the priming effect in microfluidic chambers by exposing fluorescently labelled *B. subtilis* growing on carbon-free or Carboxymethylcellulose (CMC)-containing M9-agar to different concentrations of growth medium (LB). Using time-series fluorescence microscopy, we show how the bacterial population responds to the substrate pulse over time. We derived biomass, motility, shape characteristics and spatial organization data from the images and related it to our treatments to obtain novel information on microscale processes potentially influencing the soil priming effect.

### Motility was transiently increased after the substrate pulse

*B. subtilis* is a flagellated bacterium that can swim on the agar surface and in the liquid-filled space of the chips. Our approach allowed us to distinguish stationary from moving objects on the agar patch (Fig. 2), corresponding to surface-attached single bacteria or bacterial aggregates that could not be differentiated into individual cells. Initial proportions of moving objects normalized to the agar patch area of each chip were similar across chips (average 27 ± 17.81%, Kruskal-Wallis n.s., Fig. 2 a,b), indicating similar initial conditions. Both CMC and LB concentration had significant effects on the number and proportion of moving objects, and the number of stationary objects during the time-series in the GAM (CMC - moving: p < 0.001, stationary: p = 0.018, proportion p = 0.009; LB - moving: p < 0.001; stationary: p < 0.001; proportion p = 0.04, full results provided in Table S2). A significant interactive effect of both factors was only detectable for the moving objects (p = 0.022).

**Figure 2:**
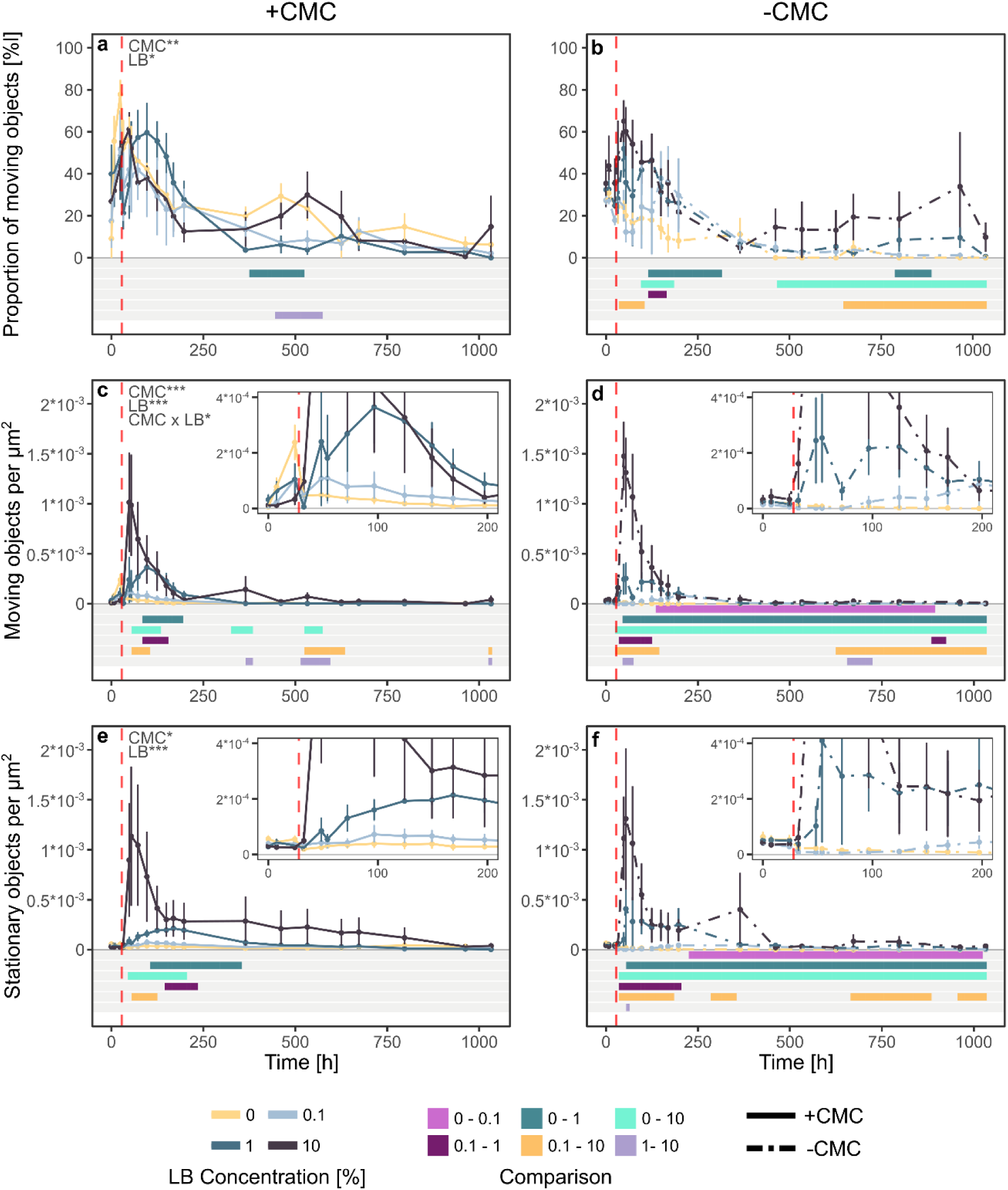
Relative proportion of moving objects in relation to total object number (**a,b**) as well as number of moving (**c,d**) and stationary objects normalized to the substrate patch area over time (**e,f**) for +CMC and -CMC chips. Depicted are mean ± SE of four chips for each timepoint and LB treatment. Plot inserts (**c-f**) represent a closeup of the first 200 hours to better distinguish differences between the lower LB groups. The red dashed line indicates the timepoint of the substrate flush. Timeseries were analyzed using generalized additive models, significant parameters of the respective model are depicted in the top left of panels **a**, **c** and **e** (p < 0.05*, p < 0.01**, p < 0.001***). Colored lanes underneath the plot indicate timespans where the respective curves were significantly different from each other in the GAM. For model details and detailed statistical results see S2.

In -CMC/0% LB, the control without any added carbon, total motility was highest at the start and decreased gradually over the course of the experiment (Fig. 2b). This pattern of a high proportion of motile objects before the flush being reduced over time afterwards was also observed when cellulose was present (+CMC/0% LB, Fig. 2a), although at overall significantly higher values (Fig. 2 a,b, Fig. S2a). The +CMC control also generally exhibited significantly more stationary and motile objects compared to the -CMC control across most of the time-series (Fig. S2a). These results suggest a baseline CMC usage by the bacteria even without LB addition, causing higher growth and motility compared to the -CMC control.

Regardless of concentration and the presence of CMC, the LB pulse increased the number of both moving and stationary objects in the short term followed by a gradual reduction over time (Fig. 2 c-f). The magnitude of the increase scaled with the pulse concentration, as also evident by the increase in the LRR relative to the -CMC/0% LB control (Fig. 3 and Fig. S3). The increase of both values was not parallel, indicated by the peak in the proportion of moving objects after the intervention in all chips that were supplied with LB (Fig. 2 a,b). Significantly different timeframes of the proportion of moving objects were rare for +CMC chips (only around 500h between 0% and 1% and 1 and 10% LB) but more common for -CMC chips, mostly between the 0% control and other LB treatments and between 0.1 and 10% LB. This pattern was also visible for the moving and stationary object counts per µm^2^ (Fig. 2 c-f), where the GAM revealed mostly short significantly different timeframes for +CMC, often directly after the pulse (Fig. 2 c,e), while differences were more sustained in the -CMC chips (Fig. 2 d,f). The presence of CMC significantly increased the response of moving and stationary objects, except for the 10% LB treatment (Fig. 3). The bacteria utilized the fresh substrate to replicate and move around on the agar surface after the pulse. Then, motility was reduced and cells became stationary. Object count was reduced over time and we could not detect a sustainably growing stationary community in the long term, however the presence of CMC generally alleviated this.

**Figure 3:**
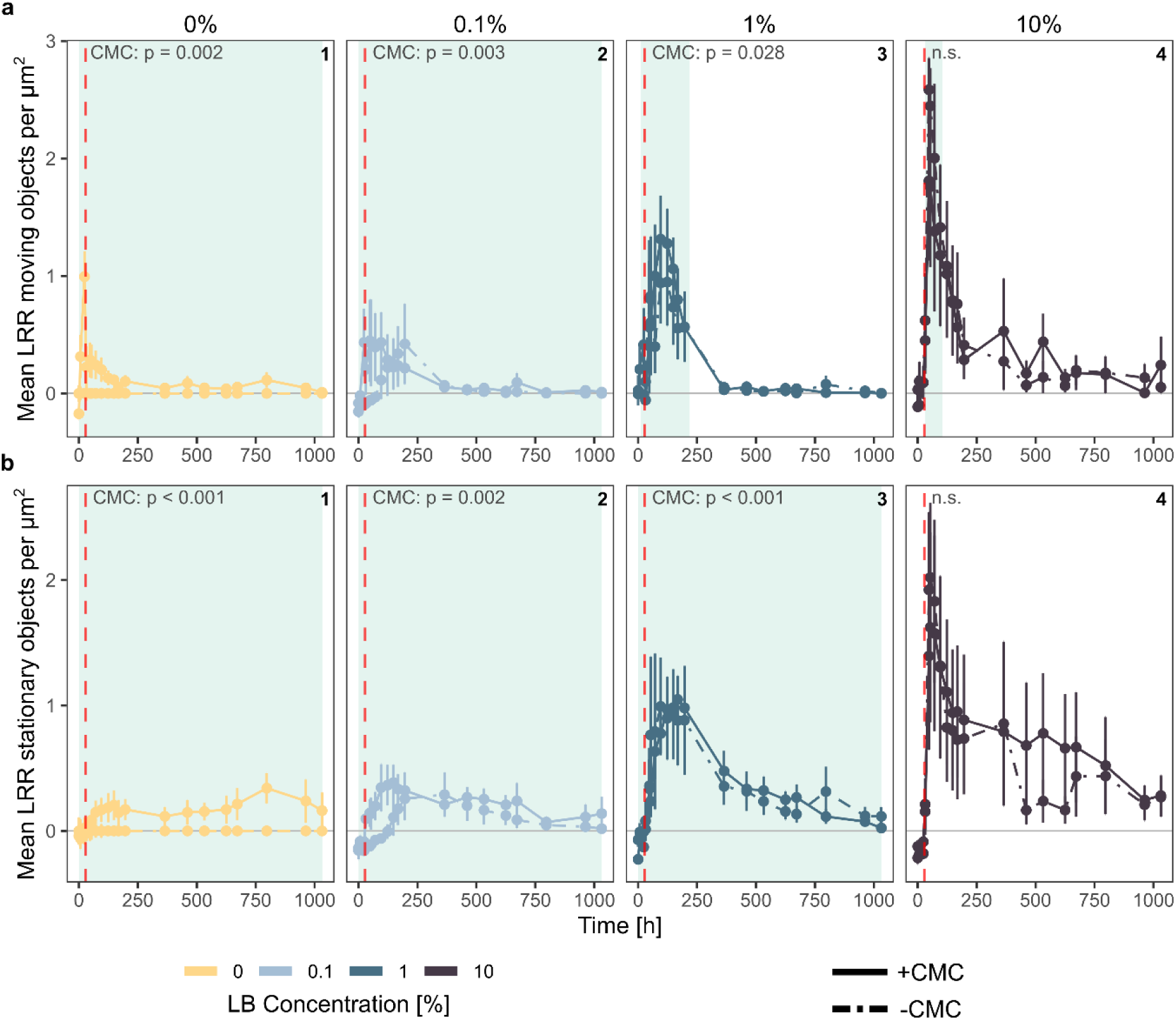
Mean log-response ratio (LRR) relative to -CMC/0% LB of moving (**a**) and stationary (**b**) object counts normalized to the patch area (mean ± SE of four chips for each timepoint and treatment). Lines compare CMC treatments across all LB concentration treatments (panels 1 to 4 each). The red dashed line indicates the timepoint of the substrate flush. Trajectories were analyzed using generalized additive models (GAM). Indicated p-values represent the effect of CMC on the respective log-response ratio in the GAMs. Significantly different timespans between +CMC and -CMC are indicated by shaded areas behind the curves. See S2 for details on model fitting procedure and detailed statistical results.

### Attached biomass responded to substrate pulse

In the following, we focused our analysis on the stationary bacteria attached to the substrate surface, as we assume these to be the ones actively decomposing CMC. The object count did not correctly represent the bacterial biomass, as larger patches which we could not resolve to single-cell level with our microscopy approach are reduced to a single object during image processing. We therefore used the area of each agarose patch that is covered by stationary fluorescent objects as a measure of attached biomass (Fig. 4). This value was significantly correlated to the stationary object count (*r*_636_ = 0.79, p < 0.001, see Fig. S4).

**Figure 4:**
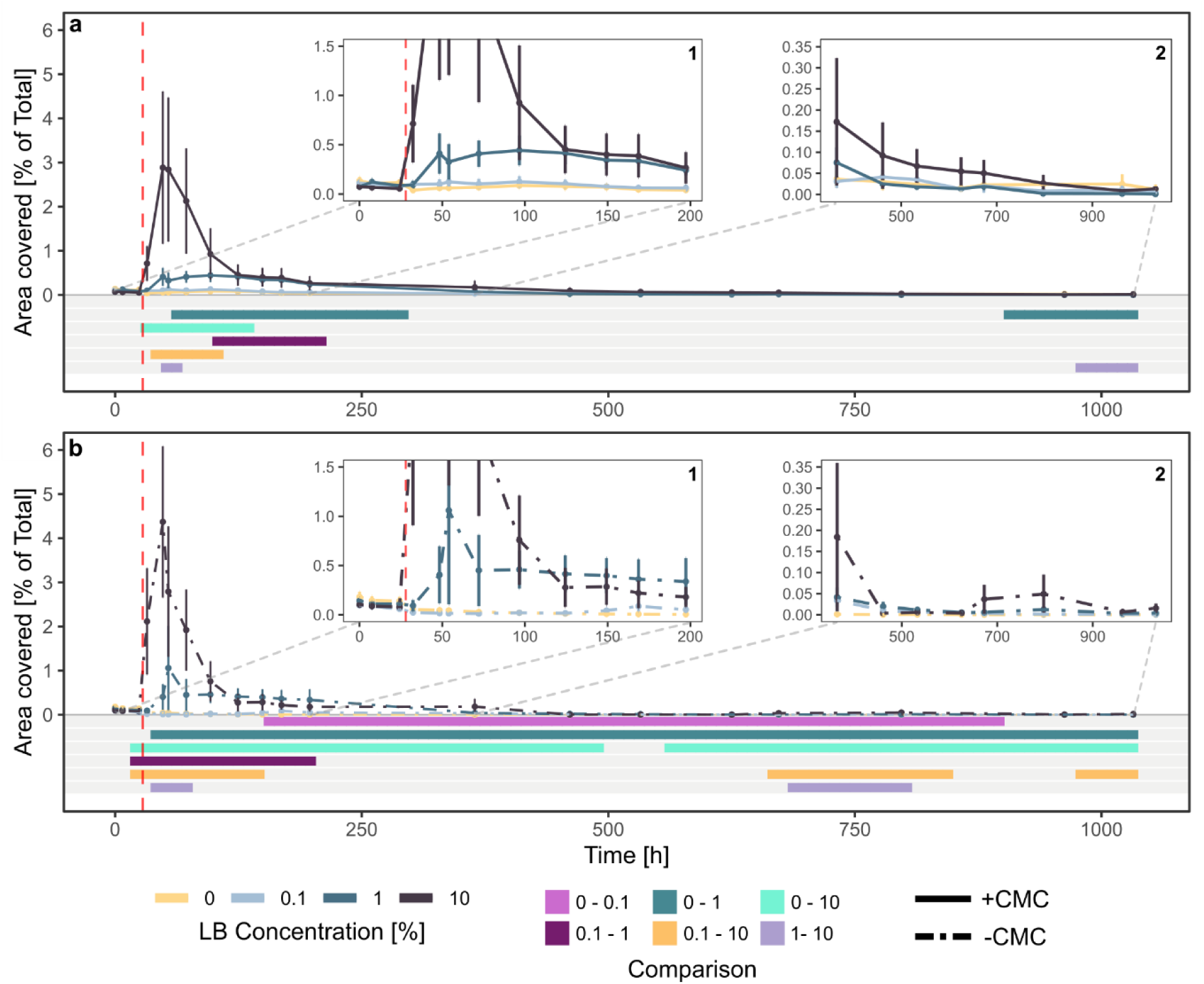
Area covered by stationary *B. subtilis* cells in % of the total CMC or M9-agar area over time for chips with (**a**) and without (**b**) CMC (mean ± SE of four chips for each timepoint and treatment). Plot inserts (1,2) are closeups of the short- and long-term responses for better visualization. The red dashed line indicates the timepoint of the substrate flush. This data can be seen as a proxy for the bacterial biomass growing on the agar, which is correlated with the total object count (see Fig. S4) but better represents larger biomass patches where individual cells could not be resolved with our approach. Timeseries were analyzed and compared using generalized additive models. Both CMC and LB concentration had significant effects on the data (CMC: p = 0.002, LB: p < 0.001, CMC x LB: n.s.), for more details see Table S2. Colored lanes underneath the plot indicate timespans where the respective curves were significantly different from each other in the GAM. For more details on model fitting procedure and analysis see S2.

When *B. subtilis* had neither complex (CMC) nor labile substrate (LB), the initial cells were flushed out by the substrate pulse with minimal media and the covered area went towards zero in the first 100-150 hours (Fig. 4 b1 and S2b). Both CMC and LB concentration significantly affected the covered area (CMC: p = 0.002; LB: p < 0.001, for full results see Table S2), with no significant interactive effect. Regardless of CMC presence, LB addition resulted in a peak of attached biomass in the short term, followed by a reduction in the long term (Fig. 4), mirroring the object count (Fig. 2). Similarly, the maximum of this peak increased with LB concentration, however there was no big difference to the 0% control at 0.1% (Fig. S5). Interestingly, only two out of the four replicates showed a strong biomass response to 10% LB (Fig. S6). Strong significant differences between time-series were again more pronounced in -CMC compared to +CMC, mostly brought on by the low attached biomass in 0% LB, underscoring that the presence of CMC had a positive effect on attached biomass. Without CMC, differences between the other LB treatments were mostly visible in the first 200 hours after the pulse and towards the end of the incubation. In the +CMC chips, trajectories of 0% and 0.1% LB chips were similar over the course of the experiment. Significant differences between other treatments again mostly appeared after the substrate pulse and towards the end of the incubation.

The LRR of the covered area relative to the no carbon control increased after the substrate pulse (Fig. 5a), scaling with LB concentration. Medium- and long-term trajectories differed from object count LRRs (Fig. 3), with generally higher values that also further increased with time. Overall, the addition of CMC increased the attached biomass compared to the chips without CMC, with significant differences of the LRRs between +CMC and -CMC for 0%, 0.1% and 1% LB. While the overall effect of CMC was again not significant in the GAM for 10% chips, their time-series still significantly differed between hour 400 to 600. The effect sizes derived from modelling further support these patterns (Fig. 5 b1, b3). LB concentration was found to increase attached biomass generally (Fig. 5 b2). When CMC was absent, this increase was exponential, however we found a hump-shaped response in the presence of CMC, with the highest effect at 1% LB, higher than its -CMC control (Fig. 5 b3). This is conversely also seen in the LRRs, where we observed a strong initial response and a steep decline followed by a gradual rebound in the long term for +CMC/10% LB (Fig. 5 a4), while the response was lower but more sustained in the medium term between hours 200 and 500 for 1% LB (Fig. 5 a3). Labile substrate addition increased the attached biomass in the chips in the short-term, but this did not translate to further growth. The response was non-linear, with intermediate LB concentrations (1%) eliciting the most sustained response after the pulse when CMC was present.

**Figure 5:**
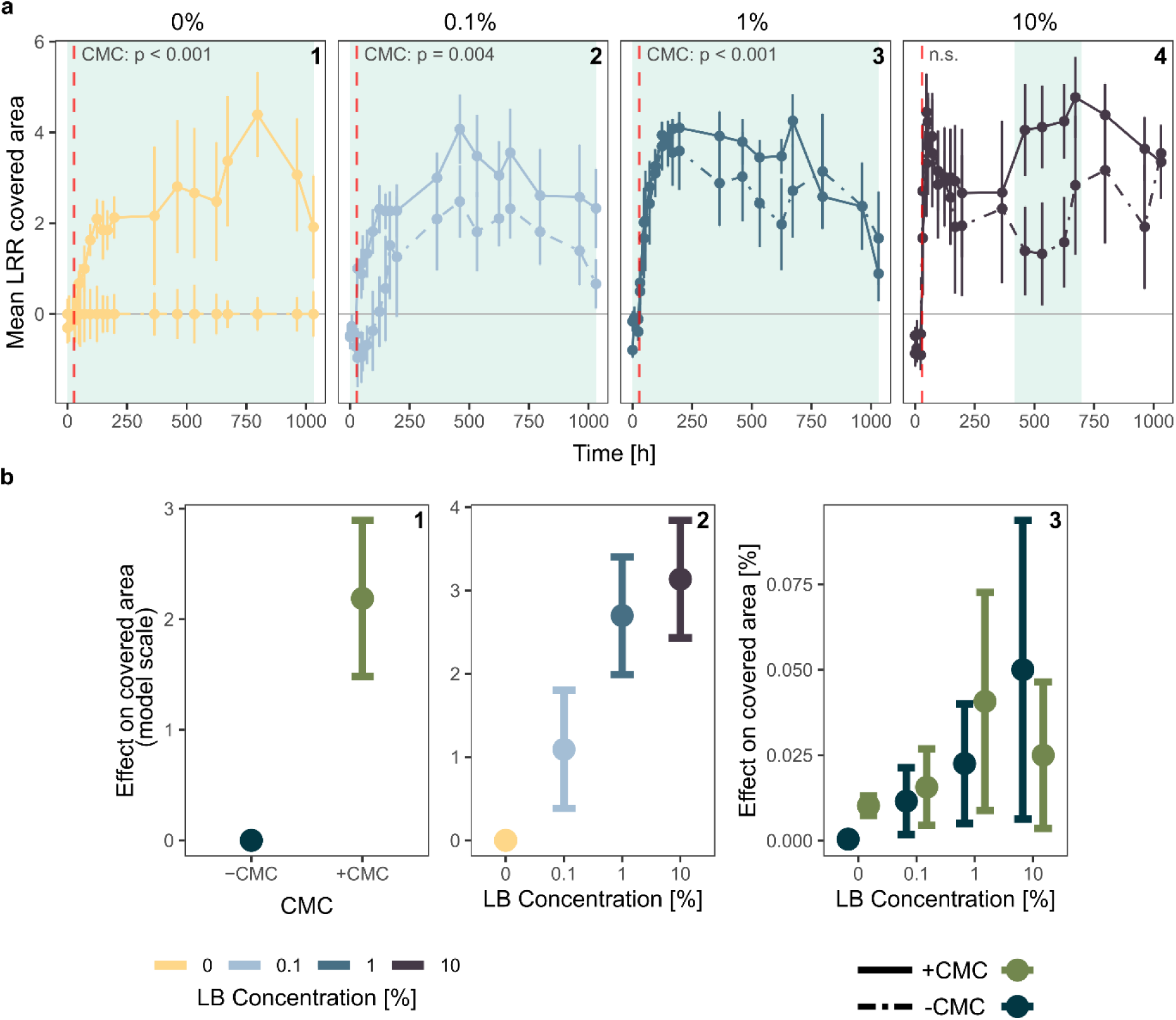
Quantification of CMC and LB treatment effects on the covered area by biomass (see Fig. 4). Mean ± SE of the time-resolved log-response ratio (LRR) of stationary attached biomass relative to the average of the -CMC/0% LB control (**a**). Lines compare CMC treatments across all LB concentration treatments (panels 1 to 4 each). The red dashed line indicates the timepoint of the substrate flush. Trajectories were analyzed using generalized additive models (GAM). Indicated p-values represent the effect of CMC on the respective LRR in the GAMs. Significantly different timespans between +CMC and -CMC are indicated by shaded areas behind the curves. Effects of experimental factors in the fitted GAM show how attached stationary biomass generally increases with CMC and LB concentration (**b**). Subplots 1 and 2 show partial effects, indicating how biomass changes with changing an isolated factor on the modelled scale (log), keeping all other model terms constant. Subplot 3 represents marginal effects, predicting the increase in biomass for each factor on the original scale, considering the full model. See S2 for details on model fitting procedure and detailed statistical results.

### Growth form affected object morphology

Generally, single *B. subtilis* cells presented as elliptical cells that elongated during growth and eventually formed chains (see Fig. 1g). High LB concentrations lead to the formation of large, irregularly shaped cell clusters that dispersed over time. This is evident by the relationship between object circularity and size (Fig. 6). Large objects with irregular shape appeared after the substrate pulse, mostly with 10% LB addition (see also Fig. S7), but then dispersed and objects became more circular again, indicating individual cells.

**Figure 6:**
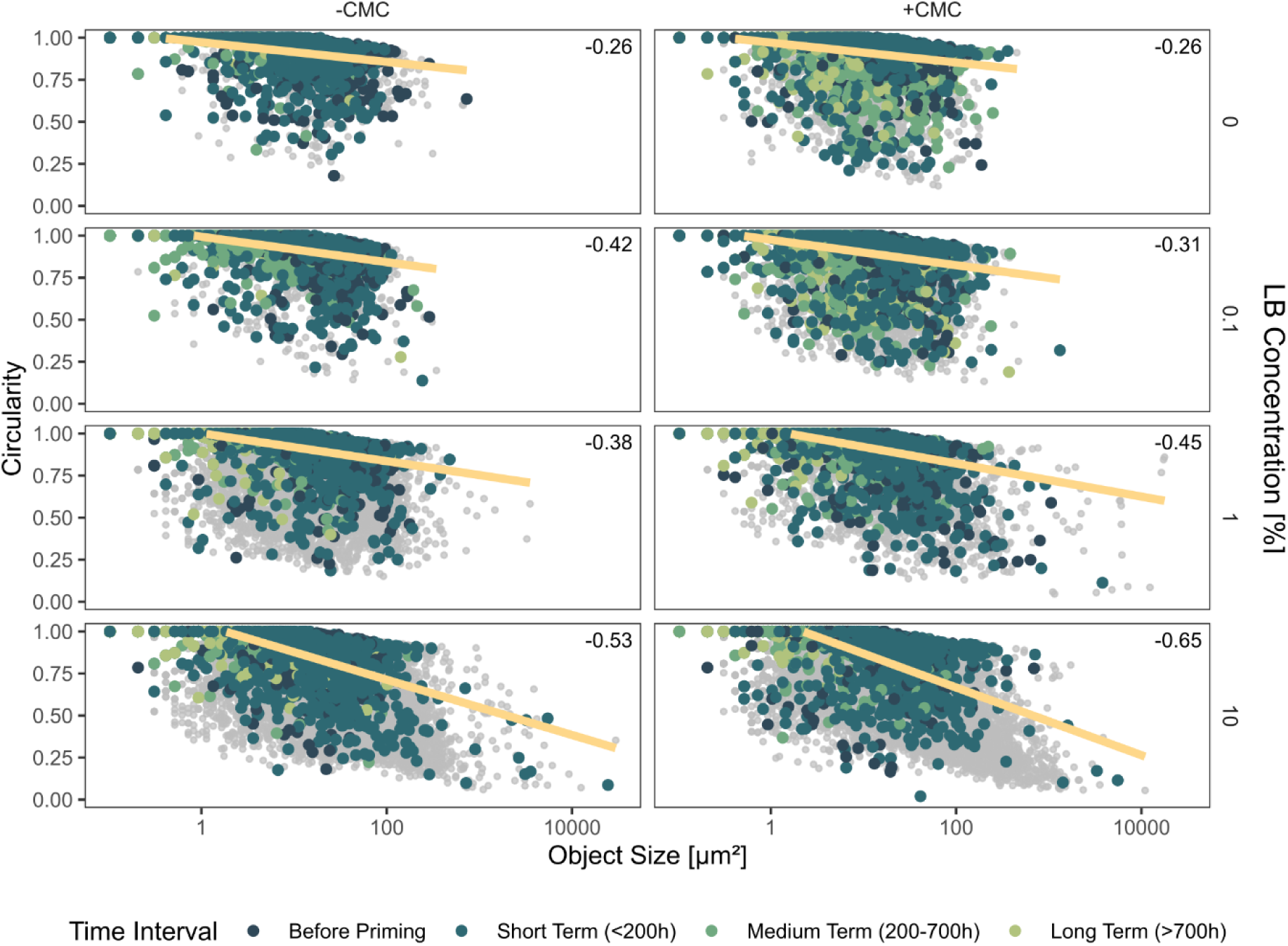
Object size related to object circularity (1 corresponds to a perfect circle) for both CMC treatments across LB concentrations for all timepoints. The yellow line indicates a linear regression fitting the data. Pearson correlation coefficients are shown in the top-right of each panel. Grey points show all detected objects (ranging from n = 4992 in –CMC 0% LB to n = 106292 in +CMC 10% LB). Colored points displayed on top are a randomly selected subset of data, as displaying all data points would be unfeasible due to their high number. They are categorized into different time intervals to visualize the evolution of the relationship across the time-series (50 points per interval).

### CMC and intermediate pulse concentrations cause clustering of bacteria

The position of objects was used to assess whether *B. subtilis* exhibited significant spatial clustering with Ripley’s K function (Fig. 1j). The deviation from complete spatial randomness (CSR) of the different replicates over time as well as their mean is visualized in a heatmap (Fig. 7a). Significant deviations were all above the CSR line, indicating clustering, with no cases of dispersion. In the low-LB treatments (0, 0.1%), CMC generally increased clustering compared to the -CMC controls. Adding LB in higher concentrations increased clustering regardless of agar substrate, especially in the short term after the priming pulse. The +CMC/1% LB treatment exhibited the most consistent clustering, underscored with the highest total significant deviation from CSR over the entire time-series (Fig. 7b). Here, we found no significant effect of LB, but a significant effect of CMC (ANOVA F_1_ = 5.528, η^2^ = 0.15, p = 0.0273). The clustering in the 10% LB treatment was reduced compared to 1%, again indicating a difference in microbial behavior between these two concentrations. The presence of a complex substrate increased the clustering of stationary cells of *B. subtilis,* which was further enhanced by intermediate substrate pulse concentrations.

**Figure 7:**
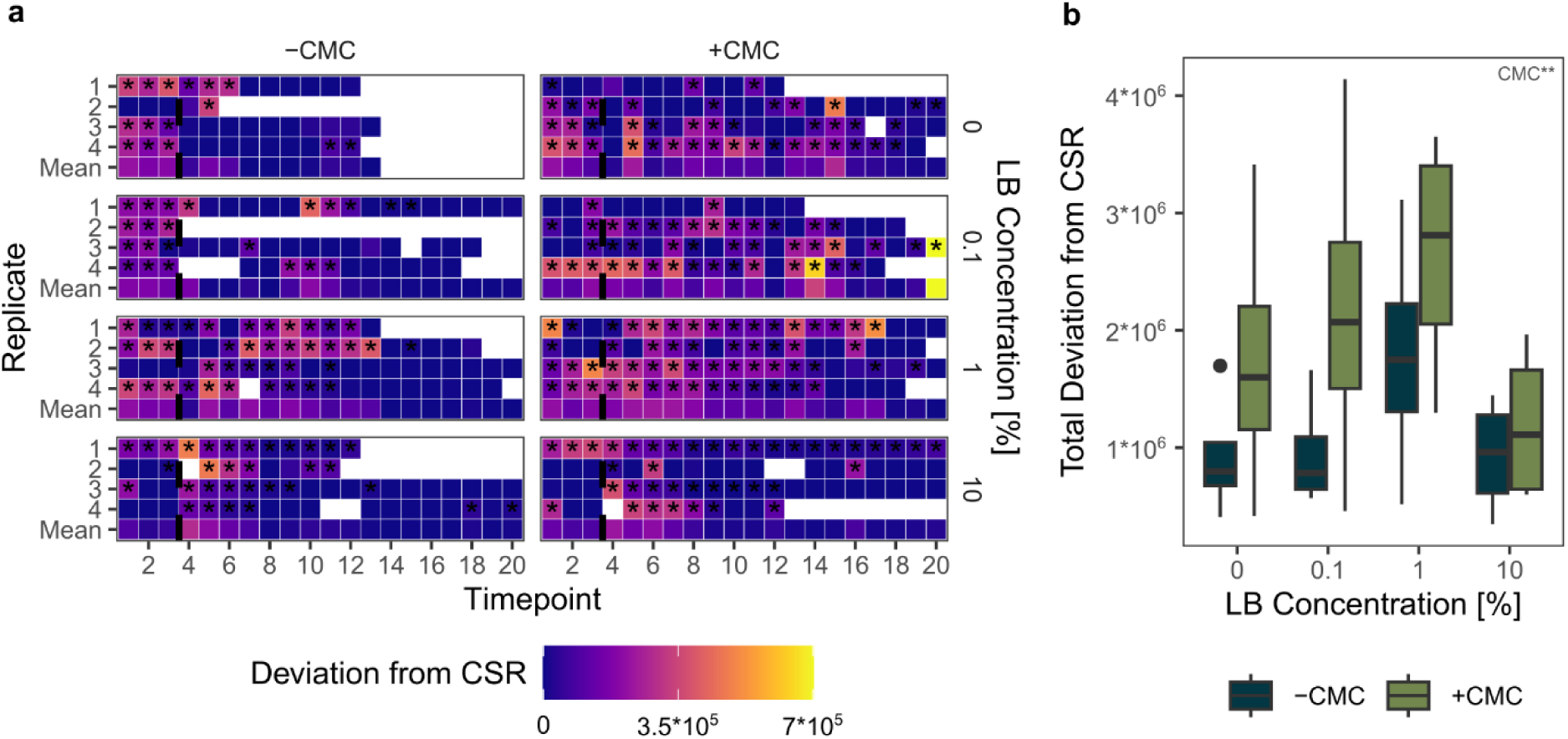
Assessment of spatial clustering of bacteria. We analyze the spatial distribution of objects in each image and calculate the deviation to complete spatial randomness (CSR) using the linearized form of Ripley’s K function. The higher this value, the higher the clustering of objects. For the individual replicates, statistical significance of this deviation was assessed using Diggle-Cressie-Loosmore-Ford tests, indicated by asterisks in the heatmap (**a**). In addition, we calculated the mean deviance for all replicates within each treatment group. The summed significant deviation from CSR over the entire time-series is shown in **b**. Significant effects of experimental factors on this value are indicated on the plot (ANOVA, p < 0.05*, p < 0.01**, p < 0.001***).

### CMC decomposition saturated between 1 and 10% LB

The amount of decomposed CMC was estimated by Congo Red staining, with lower fluorescence values indicating higher decomposition. The dye stained the carboxymethylcellulose and not some other component of the M9 agar, as indicated by the significantly higher fluorescence values when comparing all +CMC chips to all -CMC chips (Fig. S9). LB concentration significantly affected the average fluorescence intensity in the +CMC chips (Fig. 8a, Kruskal-Wallis H_4_ = 11.671, η^2^ = 0.51, p = 0.02). No treatment was significantly different from the uninoculated control, but there was a difference between the low-LB and high-LB groups. While the fluorescence of 0% and 0.1% was higher than the control, it was on average 14.29 ± 9.18% and 12.83 ± 12.05% lower in 1% and 10% respectively. This suggests lower stain binding due to CMC decomposition in the high-LB treatments.

**Figure 8:**
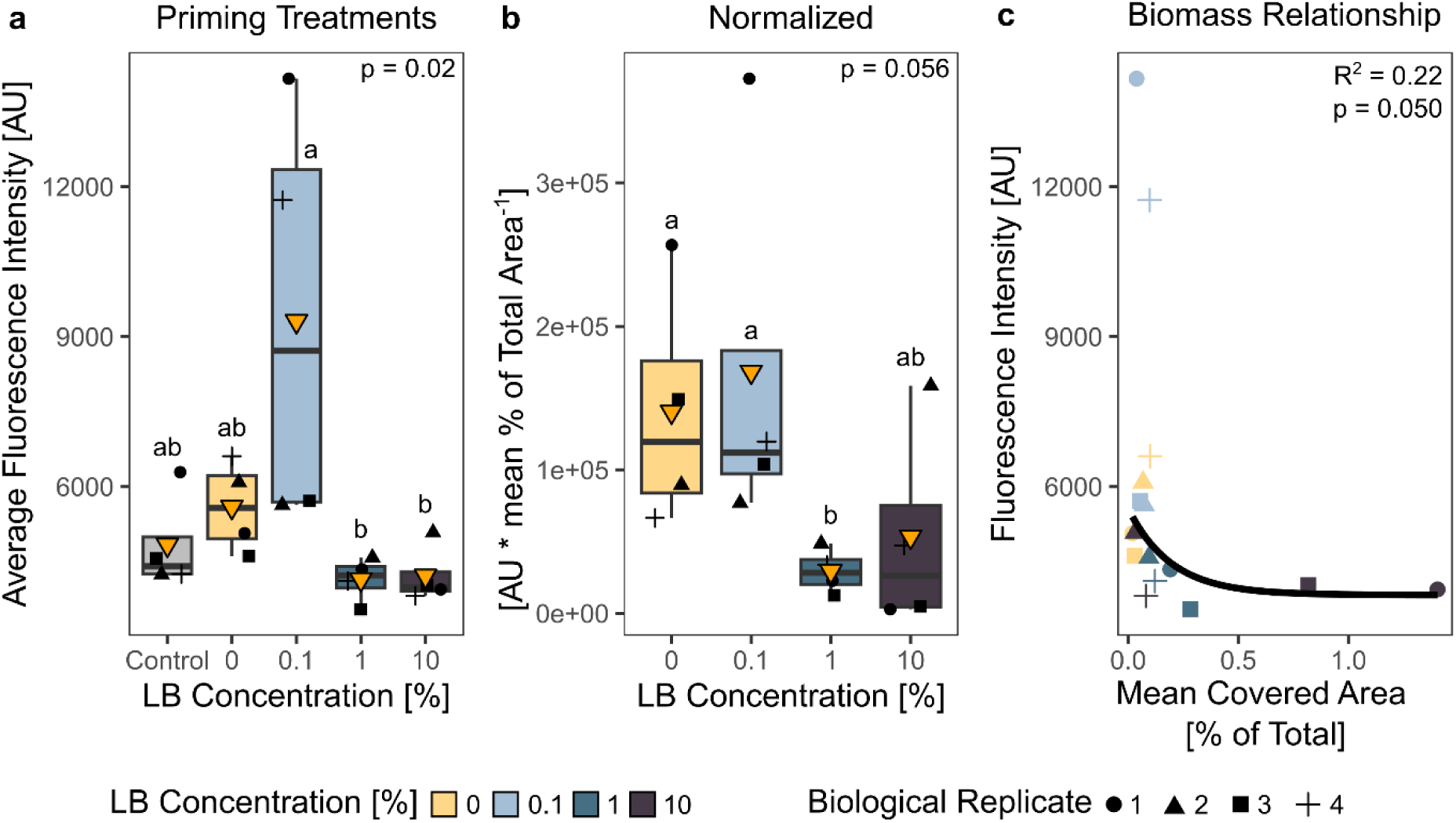
Assessment of total CMC decomposition using Congo Red staining. Average fluorescence intensity of the stained remaining substrate patches recorded at 575/590 nm excitation/emission (**a**). Lower values indicate higher decomposition. Four chips that were not inoculated by bacteria were added as control. This value was normalized to the mean attached biomass (see Fig. 4) over the entire timeseries (**b**). P-values indicate Kruskal-Wallis test results, testing for the effect of LB concentration on the raw and normalized fluorescence values. The mean attached biomass and fluorescence intensity were related by a negative exponential relationship (**c**), which is visualized here by fitting an asymptotical nonlinear least-squares regression, excluding the two high values with 0.1% LB. Here, the reported R^2^ and p-value represent a linear regression fit to the log-log transformed data.

Normalizing the average fluorescence intensity with the mean biomass (= covered area) of each chip reveals that 1% LB had the lowest mean fluorescence intensity per unit of biomass (Fig. 8b). While the difference to the low-LB chips was significant here, the overall effect of LB concentration was only marginally significant (Kruskal-Wallis H_3_ = 7.566, η^2^ = 0.38, p = 0.056). The higher mean in 10% LB mostly driven by the two replicates that did not show the same biomass response compared to the other two replicates (Fig. S6). Correlating the fluorescence to the mean attached biomass (covered area) revealed a negative exponential relationship (Fig. 8c). Measurable CMC decomposition did only happen when at least 1% LB was added, and the amount of decomposition did not scale with LB concentration but was similar in 10% LB compared to 1% LB. Additionally, decomposition was related to attached biomass, not scaling linearly with it but saturating.

### Predictors of decomposition

We used linear modelling to analyze which factors best explained CMC decomposition. A combination of different factors was tested (Table 1), collapsing highly correlated measurements into principal components. After stepwise deletion, LB concentration, average circularity and average deviation from complete spatial randomness were the best predictors for the Congo Red fluorescence (adjusted R^2^ = 0.48, F = 5.643 on 3 and 12 degrees of freedom, p = 0.012). An increase of these factors reduced the final fluorescence, indicating increased decomposition, as evident by their negative estimate (Table 1). This suggests that decomposition was strongest in the high-LB treatments and where mostly individual bacteria (high circularity) were spatially clustered (high deviation from CSR).

**Table 1:** Overview of tested linear model factors to assess their effect on CMC decomposition. Average covered area, object count per µm^2^ and maximum covered area were highly correlated (see Table S3) and collapsed into a principal component, including the first two axes to the model (PCA1 and PCA2). Model results of remaining factors after stepwise parameter deletion are shown. For more details on parameter and model selection see the supplementary material. VIF = variance inflation factor (a measure for multicollinearity).

| Final parameters | Estimate | Std. Error | t | p | Partial $R^2$ | VIF |
| --- | --- | --- | --- | --- | --- | --- |
| Intercept | 3.728 | 0.030 | 122.969 | < 0.001 |  |  |
| LB concentration | -0.860 | 0.032 | -2.649 | 0.021 | 0.37 | 1.07 |
| Average circularity | -0.010 | 0.034 | -2.945 | 0.012 | 0.42 | 1.17 |
| Average deviation from CSR | -0.068 | 0.034 | -1.986 | 0.070 | 0.25 | 1.21 |
| **Omitted parameters** |  |  |  |  |  |  |
| Average size |  |  |  |  |  |  |
| Average proportion of stationary objects |  |  |  |  |  |  |
| PCA1 |  |  |  |  |  |  |
| PCA2 |  |  |  |  |  |  |

## Discussion

We developed an experimental model system to investigate the effect of labile substrate addition on soil bacteria and how this affects long-term complex substrate degradation. A pulse of growth medium caused a transient increase in bacterial motility. The surface-attached biomass increased proportionally with pulse concentration, but this was not sustained and followed by a decrease. The presence of CMC alleviated this decrease and resulted in significant spatial clustering. Our measurements consistently showed nonlinear concentration-dependent responses in the chips with CMC, indicating saturation of decomposition at high pulse concentrations and highest spatial organization at intermediate concentration. We provide a novel and direct view into the response of soil bacteria to a substrate pulse, a first step towards understanding the microscale mechanisms behind the soil priming effect.

CMC usage by *B. subtilis* has been reported before (Stülke & Hillen, 2000) and the particular strain used here was able to utilize CMC, as evident by the higher attached and stationary biomass in the 0% and 0.1% LB treatment compared to the -CMC control. The staining method we employed was, however, not sensitive enough to capture the potential small amount of decomposition in these chips. Congo Red is usually used on larger scales and has only been used to qualitatively identify cellulolytic microorganisms (e.g. Romano et al., 2013). It can thus only give only a rough indication of complex substrate degradation. Nevertheless, as CR binding to CMC increases its fluorescence (Zemanek et al., 2017), lower fluorescence brought on by lower CMC concentration is an indication of decomposition (similar to Tyurin et al., 2021). A pre-test using purified cellulase showed strongly decreased fluorescence of the CMC agar patches (Fig. S10). When LB concentration was increased, we found evidence for higher decomposition of CMC, indicating that priming occurred in our model system. Still, decomposition measurements might be improved in future versions by employing more sophisticated methods like constant, time- and space-resolved enzymatic activity or breakdown product measurements. It might also be possible to stain the gels before the experiment and measure the decrease in fluorescence in a time-resolved manner, but this needs prior confirmation that any used stain does not infer with microbial activity.

We expected that bacterial biomass would increase with substrate pulse concentration, causing increased decomposition. In fact, the surface-attached biomass was on average highest in the 10% LB treatment shortly after the pulse, CR fluorescence indicated that CMC was decomposed in the 1% and 10% LB treatments. Studies in soil have shown that added organic matter is used up by growing microorganisms before turning to SOM (Liu et al., 2020). Similarly, the growth burst after the substrate addition was caused by the usage of the added LB medium, and not by instantly increased decomposition of CMC, as the initial response was similar between the -CMC and +CMC chips and differences mostly appeared in the medium to long term. The extra energy was likely invested into growth, not cellulase production, especially in the 10% treatment. Research has shown that *B. subtilis* can enter two different states during exponential growth. A sessile state where cells form chains and a motile state, triggered by environmental cues and allowing the population to both colonize the current habitat and move towards new ones (Colin et al., 2021; Kearns & Losick, 2005; Mukherjee & Kearns, 2014; Syvertsson et al., 2021). Since we both observed the formation of cell chains during exponential growth and an increase in relative motility, it is likely that this happened in this experiment as well. Motility is costly to a cell (Colin et al., 2021), and research into gene expression regulation in *B. subtilis* has also shown that the bacterium can face a trade-off between motility, sporulation and extracellular enzyme production (Msadek, 1999), suggesting that the decrease in relative motility over time could be caused by a shift in strategy, investing more into cellulase production in +CMC or a starvation response in the -CMC chips respectively. It is therefore possible that the observed decomposition happened in the medium- to long-term after the initial growth burst was over, which would be in line with previous observations in soil (Kuzyakov et al., 2000; Liu et al., 2020). This is further supported by the significant effect of object circularity on decomposition, indicating that coherent, irregularly shaped cell clusters are not as associated with CMC decomposition as individual, more circular cells. All in all, labile substrate addition triggering a motility response in soil bacteria has to our knowledge not been considered in the context of the priming effect. This mechanism could lead to the colonization of new habitats and a mixing of communities, possibly enhancing the degradation potential, and could therefore affect SOM decomposition. It however remains to be seen how common such a response is in soil bacteria beyond the model strain we studied.

We hypothesized that substrate addition causes some colonies to cross a density threshold where diffusion of cellulase products is no longer limiting, allowing for more efficient CMC decomposition and sustained colony growth after labile substrate is depleted. Dense cell clusters formed after the 10% pulse that could be an indication for the formation of a coherent biofilm with possible positive effects on CMC decomposition (Y. Deng & Wang, 2022), however they dispersed again in the medium-to short term and are likely more a result of the growth form of *B. subtilis* rather than coordinated biofilm formation. We therefore cannot confirm our hypothesis directly. Our results of the 1% LB treatment however still show that spatial coordination can increase the per-biomass decomposition efficiency, potentially through more efficient sharing of public goods.

In natural systems, a peak in biomass after substrate addition is often also followed by a reduction, but it usually stays elevated compared to the start or controls (e.g. Cayuela et al., 2009; Hamer & Marschner, 2005). Conversely, conditions in the chips apparently did not support sustained highly elevated biomass, even if additional C was present in the form of CMC. A small modelling exercise suggested that the population in the +CMC chips might have been close to the carrying capacity with 10% LB addition, as an asymptotical curve fit better to the (log-transformed) maximum attached biomass than a linear model (Fig. S5), so further increases in LB might not increase the peak biomass proportionally. Nutrients like Nitrogen- or Phosphorous-containing compounds in the medium could have been depleted through consumption by the high biomass, waste metabolites could have accumulated to toxic levels (L. Chen et al., 2023; Landwall & Holme, 1977) or the pH could have changed (Kram & Finkel, 2015), all possible factors preventing further growth and CMC decomposition. An investment into a moving and exploring sub-population could also have reinforced early substrate depletion as discussed above, with our spatially limited system providing no obvious benefit to the population. Interestingly, we observed a slight increase in attached biomass towards the end of the incubation in some 10% chips, possibly hinting at recycling processes, adaptation to the environment or re-colonization of the agar surface by the free-swimming population.

The addition of 0.1% LB was not enough to trigger measurable CMC decomposition in our system. Although not directly comparable, a threshold of labile substrate concentration that triggers priming has been observed in soil (e.g. Liu et al., 2017). Priming can decrease and become negative when the amount of added carbon (C) is much larger than the microbial biomass C (Blagodatskaya & Kuzyakov, 2008), or it can increase with labile substrate concentration (Di Lonardo et al., 2019; Liu et al., 2017). In our case, although we added a lot more C compared to microbial biomass C (see S13 for a back-of-the envelope calculation), CMC decomposition was concentration dependent, but it saturated between 1 and 10% LB. This could have been caused by full CMC decomposition, however the amount of C in CMC was more than double the amount that we added in 10% (about 2.93 µg C compared to 8 µg C in CMC), indicating that growth should have been much higher compared to the 0% LB controls in the case of full CMC consumption. Overall higher biomass in 10% potentially generally increased the amount of cellulases, which partially explains the LB effect on fluorescence and is in line with classical ideas of the mechanisms behind positive priming, where labile substrate-induced higher biomass leads to co-metabolization and nutrient mining (Bernard et al., 2022; Kästner et al., 2021; Zhang et al., 2025). This is supported by the fact that high decomposition was observed in the replicates with high biomass only. We do not know why growth responses differed between replicates in 10% compared to the quite consistent response in 1% (Fig. S3). They might have been possibly harvested in a physiological state where the sudden flush of nutrients caused a stress response instead of growth (Azevedo et al., 2012). With 1% LB, the peak in biomass was lower and the response was more sustained in the medium-term, which possibly prevented the auto-limitation that we observed with 10%. Here, the bacteria additionally did not need to invest in efficient CMC decomposition as they could feed on the excess LB, which is supported by the initial similarity of responses between the 10% +CMC and -CMC chips. Following the peak in biomass shortly after the priming event, the bacteria invested in coordinated CMC decomposition, clustering spatially to increase their efficiency. Although leading to roughly the same amount of decomposed substrate, the behavior of the bacteria differed and depended on added substrate concentration. If, for example, a bacterial community in an otherwise isolated micropore experiences a flush of fresh, easily available OM, higher concentration would therefore not necessarily translate to more decomposition. It is however unknown whether further decomposition would have happened if we let the experiment run for longer, the slight recovery of the attached biomass in the 10% chips towards the end could be an indication for that.

Besides such a niche case, the results in our simplified system are not directly transferable to real soil with all its complexity. Cross-feeding and successional dynamics in more diverse communities (McClure et al., 2022; Pontrelli et al., 2022) and diffusion of fresh nutrients and dilution of waste through the soil pore space (Koch, 1990; Or et al., 2007) alleviate the potential limitations that the closed system of the chip provides. Division of labor in diverse communities additionally enhances the decomposition of SOM (Giri et al., 2019), whereas our single bacterial strain seemed to not be a very efficient cellulose degrader. We were unable to show sustained growth of colonies upon reaching a density threshold, however this was likely due to limitations in our system and the chosen species and remains an interesting potential mechanism worth further exploring. Still, we show how a bacterial population reacts to a substrate pulse by creating an exploring motile sub-population and how the transiently increased biomass coordinates spatially to decompose a more complex substrate, leading to the priming effect. Different chip designs where nutrients are replenished through diffusion and utilizing more complex communities would be the logical next steps.

## Conclusion

Revealing the mechanisms behind the priming effect has been challenging as conventional bulk soil studies assess scales larger than where these processes happen. We developed a bottom-up approach to study the effect at scales relevant to soil microbes. The simplified experimental model system was able to capture some of the dynamics that have been observed at the bulk-soil scale: Growth of biomass caused by labile substrate addition, resulting in decomposition of complex substrate after the labile substrate was used up. In addition, our approach allows the characterization of spatial self-organization during decomposition and showed how bacterial motility might be affected by substrate pulses, possibly implicating SOM decomposition. Although our results are not generalizable to complex soil environments, we see this work as an initial step towards a more detailed understanding of the priming effect. In the future, improved chip designs could circumvent these limitations. The resulting modelling platform could then be further adjusted to investigate specific mechanisms, for example by implementing different substrate and nutrient conditions, tailored to the research question at hand. In addition, our experimental approach and image analysis pipeline can already be used to generally study microbial decomposition of complex organic material in a spatially and temporally resolved manner.

## Author contributions

MM and CK devised the study and experimental design. MM performed prototype development, experiments and data analysis. The first manuscript draft was written by MM, with contributions from CK. Funding was acquired by CK.

## Competing interests

The authors declare no conflicts of interest.

## Data availability

Figures and analyses presented here can be reproduced with the dataset and scripts available in a public repository at Zenodo: 10.5281/zenodo.21353091

## Acknowledgements

This work was funded by the European Research Council under the European Union’s Horizon 2020 research and innovation programme (grant agreement No 819446 to CK). The authors would like to thank Lauren V. Alteio for initially helping with the project, Marton Palatinszky and Shaul Pollak for their help and the provision of the microfluidic master wafer and *Bacillus subtilis* strain respectively, and Carlos Arrellano for his general help and advice.

## Notes

https://doi.org/10.5281/zenodo.21353092

### Competing Interest Statement

The authors have declared no competing interest.

## References

1. Aleklett, K., Kiers, E. T., Ohlsson, P., Shimizu, T. S., Caldas, V. E., & Hammer, E. C. (2018). Build your own soil: Exploring microfluidics to create microbial habitat structures. ISME Journal, 12(2), 312–319. 10.1038/ismej.2017.184

2. Arellano-Caicedo, C., Ohlsson, P., Bengtsson, M., Beech, J. P., & Hammer, E. C. (2021). Habitat geometry in artificial microstructure affects bacterial and fungal growth, interactions, and substrate degradation. Communications Biology, 4(1). 10.1038/s42003-021-02736-4

3. Azevedo, N. F., Bragança, S. M., Simões, L. C., Cerqueira, L., Almeida, C., Keevil, C. W., & Vieira, M. J. (2012). Proposal for a method to estimate nutrient shock effects in bacteria. BMC Research Notes, 5(1), 422. 10.1186/1756-0500-5-422

4. Baddeley, A., Diggle, P. J., Hardegen, A., Lawrence, T., Milne, R. K., & Nair, G. (2014). On tests of spatial pattern based on simulation envelopes. Ecological Monographs, 84(3), 477–489. 10.1890/13-2042.1

5. Baddeley, A., Rubak, E., & Turner, R. (2015). Spatial Point Patterns: Methodology and Applications with R. Chapman and Hall/CRC.

6. Bernard, L., Basile-Doelsch, I., Derrien, D., Fanin, N., Fontaine, S., Guenet, B., Karimi, B., Marsden, C., & Maron, P. A. (2022). Advancing the mechanistic understanding of the priming effect on soil organic matter mineralisation. Functional Ecology, 36(6), 1355–1377. 10.1111/1365-2435.14038

7. Blagodatskaya, E., & Kuzyakov, Y. (2008). Mechanisms of real and apparent priming effects and their dependence on soil microbial biomass and community structure: Critical review. Biology and Fertility of Soils, 45(2), 115–131. 10.1007/s00374-008-0334-y

8. Cayuela, M. L., Sinicco, T., & Mondini, C. (2009). Mineralization dynamics and biochemical properties during initial decomposition of plant and animal residues in soil. Applied Soil Ecology, 41(1), 118–127. 10.1016/j.apsoil.2008.10.001

9. Chen, L., Wang, C., & Su, J. (2023). Understanding the Effect of Different Glucose Concentrations in the Oligotrophic Bacterium Bacillus subtilis BS-G1 through Transcriptomics Analysis. Microorganisms, 11(10), 2401. 10.3390/microorganisms11102401

10. Chen, R., Senbayram, M., Blagodatsky, S., Myachina, O., Dittert, K., Lin, X., Blagodatskaya, E., & Kuzyakov, Y. (2014). Soil C and N availability determine the priming effect: Microbial N mining and stoichiometric decomposition theories. Global Change Biology, 20(7), 2356–2367. 10.1111/gcb.12475

11. Colin, R., Ni, B., Laganenka, L., & Sourjik, V. (2021). Multiple functions of flagellar motility and chemotaxis in bacterial physiology. FEMS Microbiology Reviews, 45(6), fuab038. 10.1093/femsre/fuab038

12. Deng, Y., & Wang, S. Y. (2022). Sorption of Cellulases in Biofilm Enhances Cellulose Degradation by Bacillus subtilis. Microorganisms, 10(8), 1505. 10.3390/microorganisms10081505

13. Deng, Y.-J., & Wang, S. Y. (2016). Synergistic growth in bacteria depends on substrate complexity. Journal of Microbiology, 54(1), 23–30. 10.1007/s12275-016-5461-9

14. Di Lonardo, D. P., De Boer, W., Zweers, H., & Van Der Wal, A. (2019). Effect of the amount of organic trigger compounds, nitrogen and soil microbial biomass on the magnitude of priming of soil organic matter. PLOS ONE, 14(5), e0216730. 10.1371/journal.pone.0216730

15. D’Souza, G. G., Povolo, V. R., Keegstra, J. M., Stocker, R., & Ackermann, M. (2021). Nutrient complexity triggers transitions between solitary and colonial growth in bacterial populations. The ISME Journal, 15(9), 2614–2626. 10.1038/s41396-021-00953-7

16. Ebrahimi, A., Schwartzman, J., & Cordero, O. X. (2019). Cooperation and spatial self-organization determine rate and efficiency of particulate organic matter degradation in marine bacteria. Proceedings of the National Academy of Sciences, 116(46), 23309–23316. 10.1073/pnas.1908512116

17. Ershov, D., Phan, M.-S., Pylvänäinen, J. W., Rigaud, S. U., Le Blanc, L., Charles-Orszag, A., Conway, J. R. W., Laine, R. F., Roy, N. H., Bonazzi, D., Duménil, G., Jacquemet, G., & Tinevez, J.-Y. (2022). TrackMate 7: Integrating state-of-the-art segmentation algorithms into tracking pipelines. Nature Methods, 19(7), 829–832. 10.1038/s41592-022-01507-1

18. Giri, S., Waschina, S., Kaleta, C., & Kost, C. (2019). Defining Division of Labor in Microbial Communities. Journal of Molecular Biology, 431(23), 4712–4731. 10.1016/j.jmb.2019.06.023

19. Guseva, K., Mohrlok, M., Alteio, L., Schmidt, H., Pollak, S., & Kaiser, C. (2024). Bacteria face trade-offs in the decomposition of complex biopolymers. PLOS Computational Biology, 20(8), e1012320. 10.1371/journal.pcbi.1012320

20. Hamer, U., & Marschner, B. (2005). Priming effects in soils after combined and repeated substrate additions. Geoderma, 128(1–2), 38–51. 10.1016/j.geoderma.2004.12.014

21. Kaiser, C., Dieckmann, U., & Franklin, O. (2017). The rhizosphere priming effect explained by microscale interactions among enzyme producing microbes [Conference Presentation]. *Geophysical Research Abstracts*, EGU General Assembly 2017, 19(EGU2017-19214).

22. Kästner, M., Miltner, A., Thiele-Bruhn, S., & Liang, C. (2021). Microbial Necromass in Soils—Linking Microbes to Soil Processes and Carbon Turnover. Frontiers in Environmental Science, 9. 10.3389/fenvs.2021.756378

23. Kearns, D. B., & Losick, R. (2005). Cell population heterogeneity during growth of *Bacillus subtilis*. Genes & Development, 19(24), 3083–3094. 10.1101/gad.1373905

24. Koch, A. L. (1990). Diffusion The Crucial Process in Many Aspects of the Biology of Bacteria. In K. C. Marshall (Hrsg.), Advances in Microbial Ecology (S. 37–70). Springer US. 10.1007/978-1-4684-7612-5_2

25. Kram, K. E., & Finkel, S. E. (2015). Rich Medium Composition Affects Escherichia coli Survival, Glycation, and Mutation Frequency during Long-Term Batch Culture. Applied and Environmental Microbiology, 81(13), 4442–4450. 10.1128/AEM.00722-15

26. Kuzyakov, Y. (2010). Priming effects: Interactions between living and dead organic matter. Soil Biology and Biochemistry, 42(9), 1363–1371. 10.1016/j.soilbio.2010.04.003

27. Kuzyakov, Y., Friedel, J. K., & Stahr, K. (2000). Review of mechanisms and quantification of priming effects. Soil Biology & Biochemistry, 32, 1485–1498.

28. Lal, R., & Stewart, B. A. (2019). Soil and Climate. CRC Press.

29. Landwall, P., & Holme, T. (1977). Removal of Inhibitors of Bacterial Growth by Dialysis Culture. Journal of General Microbiology, 103(2), 345–352. 10.1099/00221287-103-2-345

30. Lehmann, J., Hansel, C. M., Kaiser, C., Kleber, M., Maher, K., Manzoni, S., Nunan, N., Reichstein, M., Schimel, J. P., Torn, M. S., Wieder, W. R., & Kögel-Knabner, I. (2020). Persistence of soil organic carbon caused by functional complexity. Nature Geoscience, 13(8), 529–534. 10.1038/s41561-020-0612-3

31. Liu, X.-J. A., Finley, B. K., Mau, R. L., Schwartz, E., Dijkstra, P., Bowker, M. A., & Hungate, B. A. (2020). The soil priming effect: Consistent across ecosystems, elusive mechanisms. Soil Biology and Biochemistry, 140, 107617. 10.1016/j.soilbio.2019.107617

32. Liu, X.-J. A., Sun, J., Mau, R. L., Finley, B. K., Compson, Z. G., Van Gestel, N., Brown, J. R., Schwartz, E., Dijkstra, P., & Hungate, B. A. (2017). Labile carbon input determines the direction and magnitude of the priming effect. Applied Soil Ecology, 109, 7–13. 10.1016/j.apsoil.2016.10.002

33. McClure, R., Farris, Y., Danczak, R., Nelson, W., Song, H.-S., Kessell, A., Lee, J.-Y., Couvillion, S., Henry, C., Jansson, J. K., & Hofmockel, K. S. (2022). Interaction Networks Are Driven by Community-Responsive Phenotypes in a Chitin-Degrading Consortium of Soil Microbes. mSystems, 7(5), 1–21. 10.1128/msystems.00372-22

34. Msadek, T. (1999). When the going gets tough: Survival strategies and environmental signaling networks in Bacillus subtilis. 7(5).

35. Mukherjee, S., & Kearns, D. B. (2014). The Structure and Regulation of Flagella in *Bacillus subtilis*. Annual Review of Genetics, 48(1), 319–340. 10.1146/annurev-genet-120213-092406

36. Or, D., Smets, B. F., Wraith, J. M., Dechesne, A., & Friedman, S. P. (2007). Physical constraints affecting bacterial habitats and activity in unsaturated porous media – a review. Advances in Water Resources, 30(6–7), 1505–1527. 10.1016/j.advwatres.2006.05.025

37. Pontrelli, S., Szabo, R., Pollak, S., Schwartzman, J., Ledezma-Tejeida, D., Cordero, O. X., & Sauer, U. (2022). Metabolic crossfeeding structures the assembly of polysaccharide degrading communities. Science Advances, 8(February), 1–11. 10.1126/sciadv.abk3076

38. R Core Team. (2025). R: A Language and Environment for Statistical Computing [Software]. R Foundation for Statistical Computing.

39. Raynaud, X., & Nunan, N. (2014). Spatial ecology of bacteria at the microscale in soil. PLoS ONE, 9(1). 10.1371/journal.pone.0087217

40. Ripley, B. D. (1988). Statistical Inference for Spatial Processes. Cambridge University Press.

41. Romano, N., Gioffré, A., Sede, S. M., Campos, E., Cataldi, A., & Talia, P. (2013). Characterization of Cellulolytic Activities of Environmental Bacterial Consortia from an Argentinian Native Forest. Current Microbiology, 67(2), 138–147. 10.1007/s00284-013-0345-2

42. Rusconi, R., Garren, M., & Stocker, R. (2014). Microfluidics expanding the frontiers of microbial ecology. Annual Review of Biophysics, 43(1), 65–91. 10.1146/annurev-biophys-051013-022916

43. Sharma, K., Palatinszky, M., Nikolov, G., Berry, D., & Shank, E. A. (2020). Transparent soil microcosms for live-cell imaging and non-destructive stable isotope probing of soil microorganisms. eLife, 9, 1–28. 10.7554/eLife.56275

44. Sharma, R., Salwan, R., Kaur, R., & Sharma, V. (2025). Microbial contributions to soil carbon sequestration. In Advances in Botanical Research (Bd. 116, S. 115–130). Elsevier. 10.1016/bs.abr.2025.04.002

45. Souza, C. P., Burbano-Rosero, E. M., Almeida, B. C., Martins, G. G., Albertini, L. S., & Rivera, I. N. G. (2009). Culture medium for isolating chitinolytic bacteria from seawater and plankton. World Journal of Microbiology and Biotechnology, 25(11), 2079–2082. 10.1007/s11274-009-0098-z

46. Stülke, J., & Hillen, W. (2000). Regulation of Carbon Catabolism in Bacillus Species. Annual Review of Microbiology, 54(1), 849–880. 10.1146/annurev.micro.54.1.849

47. Syvertsson, S., Wang, B., Staal, J., Gao, Y., Kort, R., & Hamoen, L. W. (2021). Different Resource Allocation in a Bacillus subtilis Population Displaying Bimodal Motility. Journal of Bacteriology, 203(12). 10.1128/JB.00037-21

48. Tyurin, A. A., Suhorukova, A. V., Deineko, I. V., Pavlenko, O. S., Fridman, V. A., & Goldenkova-Pavlova, I. V. (2021). A high throughput assay of lichenase activity with Congo red dye in plants. Plant Methods, 17(1), 102. 10.1186/s13007-021-00801-x

49. Wood, S. (2017). *Generalized Additive Models: An Introduction with R* (2. Aufl.). Chapman and Hall/CRC.

50. Yu, Z., Chen, L., Pan, S., Li, Y., Kuzyakov, Y., Xu, J., Brookes, P. C., & Luo, Y. (2018). Feedstock determines biochar-induced soil priming effects by stimulating the activity of specific microorganisms. European Journal of Soil Science, 69(3), 521–534. 10.1111/ejss.12542

51. Zemanek, G., Jagusiak, A., Chłopaś, K., Piekarska, B., & Stopa, B. (2017). Congo red fluorescence upon binding to macromolecules – a possible explanation for the enhanced intensity. Bio-Algorithms and Med-Systems, 13(2), 69–78. 10.1515/bams-2017-0010

52. Zhang, S., Zhang, Z., Wang, F., Huang, X., Chen, X., Wang, Y., Li, C., & Li, H. (2025). Advancing the comprehensive understanding of soil organic carbon priming effect: Definitions, mechanisms, influencing factors, and future perspectives. Environmental Geochemistry and Health, 47(6), 201. 10.1007/s10653-025-02516-7

53. Zhu, X., Wang, K., Yan, H., Liu, C., Zhu, X., & Chen, B. (2022). Microfluidics as an Emerging Platform for Exploring Soil Environmental Processes: A Critical Review. Environmental Science & Technology, 56(2), 711–731. 10.1021/acs.est.1c03899

