## Supplementary Material for "An experimental model system to investigate microscale mechanisms behind the soil priming effect"

#### S1: Example pictures

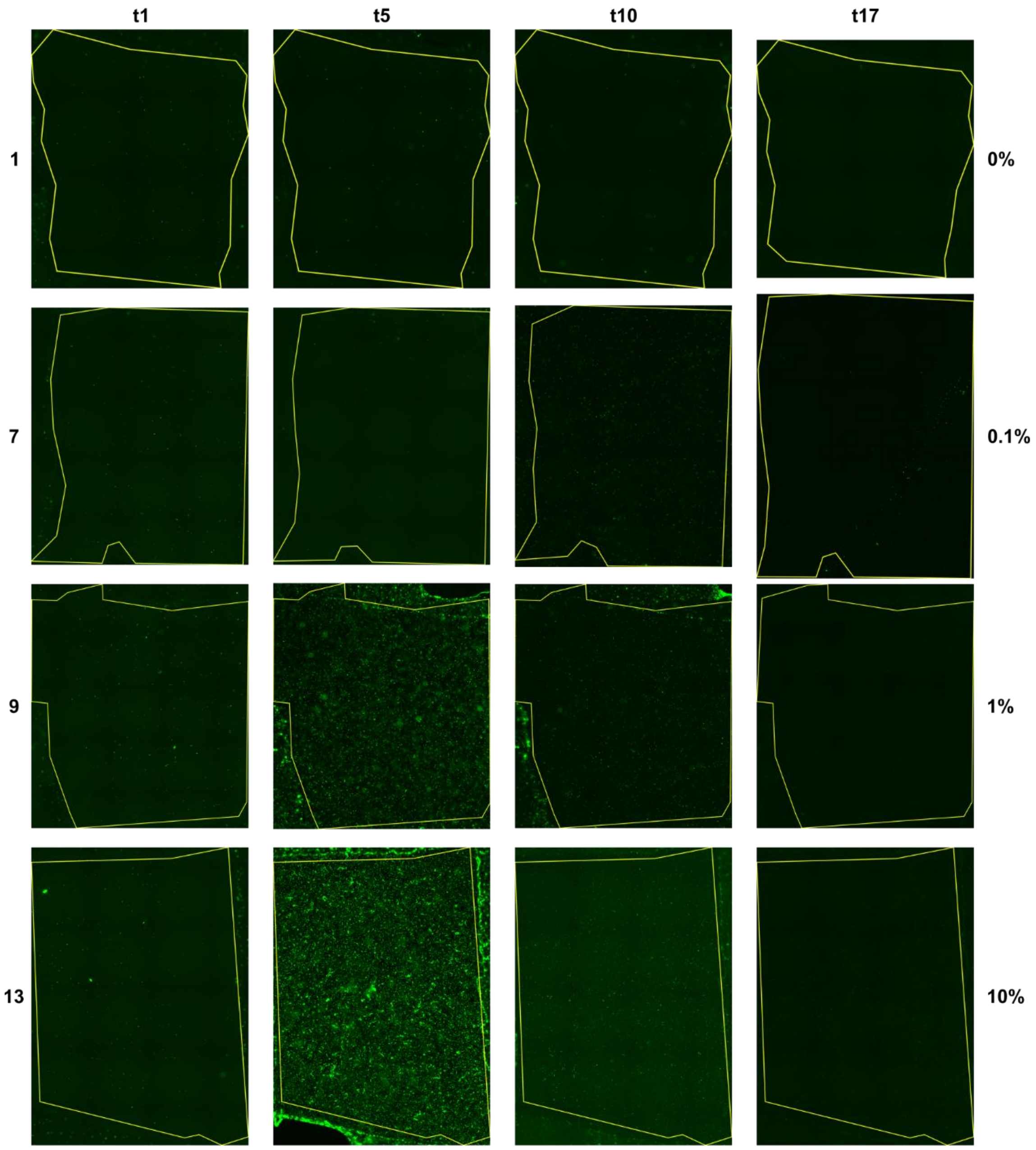

**Figure S1:** Images of +CMC chips (Chips 1, 7, 9, 13) at 4 different timepoints, for four different LB concentrations to illustrate the results exemplary. Images were converted to 8 bit and saved as portable network graphics (png) for this purpose. No further processing was done on these images. The yellow outline represents the agar patch area, which becomes visible when increasing the contrast. The complete dataset with higher resolution ".tif" files and image processing results is additionally provided. Original .lif files are available upon request.

**S2: Generalized Additive Models**

Time-series were analzyed using function gam() of R package “mgcv” (1.9-1, Wood, 2017). The used model formula was:

Reponse ~ CMC * LB Concentration + s(Time, by =CMC, k = […], bs = “cr”) + s(Chip, bs = “re”)

with cubic regression splines for time and a random spline for each chip. The number of knots k was chosen so that the effective degrees of freedom (edf) were lower than k’, and the k-index was close to 1 and not significant (checked with function check.gam()). Different transformations and model families were tested for each dataset. The best performing models were chosen based on visual inspection of model diagnostics (using function appraise() of R package “gratia” (0.11.1, Simpson, 2024) as well as their adjusted $R^2$ values and Akaike Information Criterion (AIC). Biomass data as well as the normalized number of moving and stationary objects was modelled assuming an underlying Tweedie distribution and a log-link function, which produced the best results as the data was right-skewed and contained zeroes. For the proportion of motile objects, the response was $\log_{10}$+1 transformed and modelled using an underlying gaussian distribution. Time-series of log-response ratios of normalized moving and stationary object counts (Fig. 2) and covered area by stationary objects (Fig. 5a) were analyzed with a similar GAM, but for each LB concentration level separately with the formula:

Reponse ~ CMC + s(Time, by = interaction(CMC, LB Concentration), k = […], bs = “cr”) + s(Chip, bs = “re”)

We computed significantly different timespans between model predictions using function plot_diff() of package “itsadug” (2.4.1, van Rij et al., (2022). Significantly different timespans (where the confidence interval of the difference smooth excluded 0) between LB treatments are shown as colored lanes under the trajectories in Figure 2 and Figure 4 or as shaded areas when comparing CMC treatments (Fig. 3, 5a, S2a,b). Partial effects (effect of one parametric factor on the response when keeping all other model factors constant) and marginal effects (effect of parametric factor on response considering the full model) shown in Figure 5 were obtained via functions draw() of “gratia” and plot_predictions() of package “marginaleffects” (Arel-Bundock et al., 2024) respectively. Detailed model properties and test results of parametric model terms are provided in tables S1 and S2 below respectively.

**Table S1: Model properties**

| Variable | | Adj. $R^2$ | n | Family | Link |
|---|---|---|---|---|---|
| Moving objects | | 0.682 | 638 | Tweedie | log |
| Stationary objects | | 0.764 | 638 | Tweedie | log |
| Proportion of moving objects | | 0.613 | 613 | Gaussian | identity |
| Covered area | | 0.753 | 638 | Tweedie | log |
| Log-response ratios | **LB level** | | | | |
| LRR moving objects | 0 | 0.252 | 160 | Gaussian | identity |
| | 01 | 0.166 | 159 | | |
| | 1 | 0.381 | 159 | | |
| | 10 | 0.410 | 160 | | |
| LRR stationary objects | 0 | 0.353 | 160 | | |
| | 0.1 | 0.423 | 159 | | |
| | 1 | 0.513 | 159 | | |
| | 10 | 0.152 | 160 | | |
| LRR covered area | 0 | 0.542 | 160 | | |
| | 0.1 | 0.516 | 159 | | |
| | 1 | 0.621 | 159 | | |
| | 10 | 0.228 | 160 | | |

**Table S2: Significance of parametric model terms**

| Variable | | Factor | df | F | p |
|---|---|---|---|---|---|
| Moving objects | | CMC | 1 | 13.677 | <0.001 |
| | | LB concentration | 3 | 9.764 | <0.001 |
| | | CMC x LB concentration | 3 | 3.241 | 0.022 |
| Stationary objects | | CMC | 1 | 5.679 | 0.018 |
| | | LB concentration | 3 | 6.605 | <0.001 |
| | | CMC x LB concentration | 3 | 1.342 | 0.26 |
| Proportion of moving objects | | CMC | 1 | 6.884 | 0.008 |
| | | LB concentration | 3 | 2.78 | 0.04 |
| | | CMC x LB concentration | 3 | 1.621 | 0.183 |
| Covered area | | CMC | 1 | 9.612 | 0.002 |
| | | LB concentration | 3 | 8.432 | <0.001 |
| | | CMC x LB concentration | 3 | 2.016 | 0.111 |
| | **LB level** | | | **t** | **p** |
| LRR moving objects | 0 | CMC | 1 | 3.082 | 0.002 |
| | 0.1 | | | 2.994 | 0.003 |
| | 1 | | | 2.219 | 0.028 |
| | 10 | | | -1.791 | 0.075 |
| LRR stationary objects | 0 | | | 5.412 | <0.001 |
| | 0.1 | | | 3.180 | 0.002 |
| | 1 | | | 4.663 | <0.001 |
| | 10 | | | -1.388 | 0167 |
| LRR covered area | 0 | | | 6.616 | <0.001 |
| | 0.1 | | | 2.903 | 0.004 |
| | 1 | | | 5.496 | <0.001 |
| | 10 | | | 0.160 | 0.111 |

**S3: CMC comparisons**

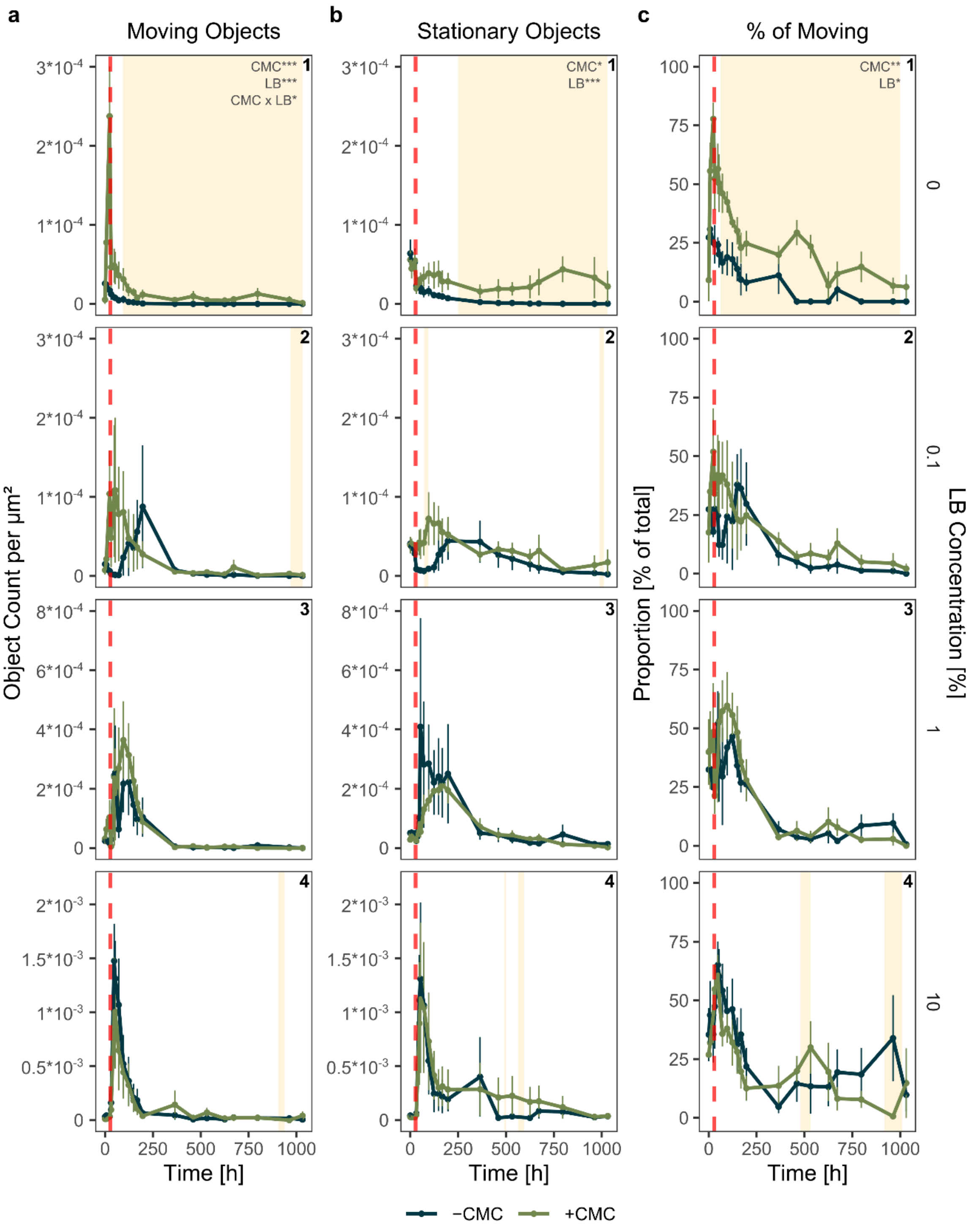

**Figure S2a:** Total number of moving and stationary objects normalized to the substrate patch area over time (a,b) as well as the relative proportion of moving objects (c). Data corresponds to Fig. 2 but is contrasted by CMC levels. Depicted are mean ± SE of four chips for each timepoint and treatment. Time-series were analyzed using generalized additive models. Significant parametric model terms are indicated in the top right of the first panel of each subplot (p < 0.05*, p < 0.01**, p < 0.001***). Shaded areas indicate timespans where the two curves (+ and - CMC) are significantly different from each other. The red dashed line indicates the timepoint of the substrate flush.

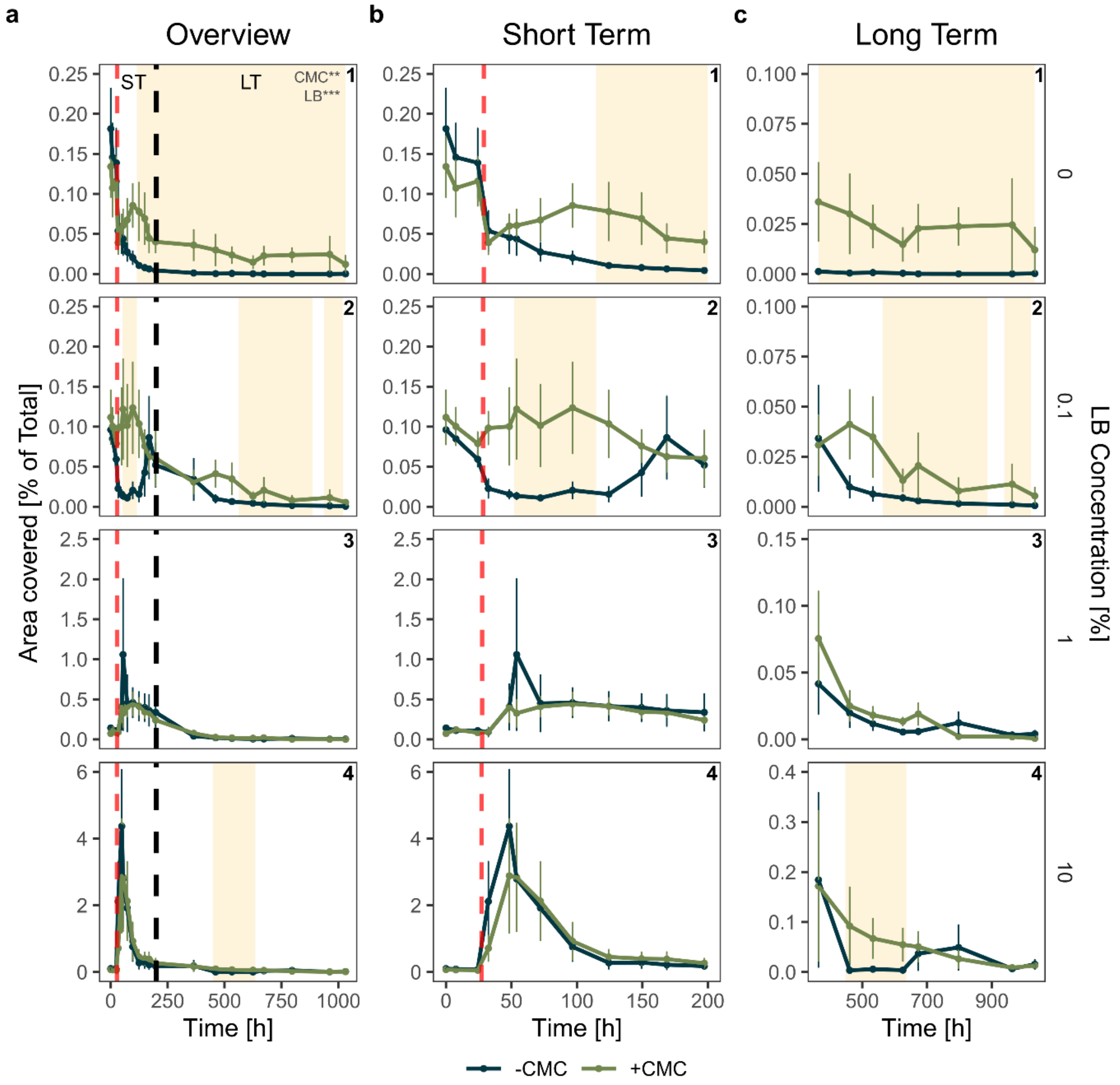

**Figure S2b:** Area covered by stationary *B. subtilis* cells in % of the total CMC or M9-agar area over time for the different priming treatments (mean ± SE of four chips for each timepoint and treatment). Data corresponds to Fig. 3 but is contrasted by CMC levels. An overview is provided in **a**, short-term and long term responses are shown with different y-axis scaling in **b** and **c** respectively for visualization purposes. The black dashed line in **a** indicates where the plots were split. This data can be seen as a proxy for the bacterial biomass growing on the agar, which is correlated with the total object count (see Fig. S4) but better represents larger biomass patches where individual cells could not be resolved with our approach. Time-series were analyzed and compared using generalized additive models.

Significant parametric model terms are indicated in the top right of the first panel (p < 0.05*, p < 0.01**, p < 0.001***). Significantly different timespans between + and -CMC are indicated with the shaded areas. The red dashed line indicates the timepoint of the substrate flush.

**S4: Maximum object count**

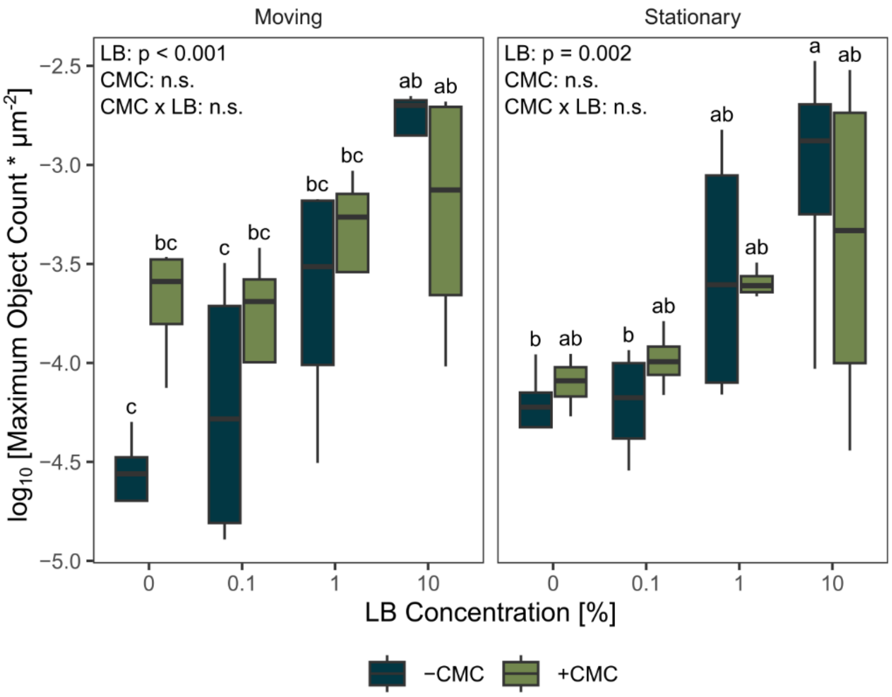

**Figure S3:** Boxplots (median and interquartile range) showing the maximum object count per LB concentration and CMC treatment for motile and stationary objects on a $\log_{10}$-$\log_{10}$ scale. A two-way ANOVA was performed for each movement type, p-values are indicated in the Figure (Moving – LB: $F_3 = 7.861$; Stationary – LB: $F_3 = 6.651$, $\alpha = 0.05$). Lowercase letters represent the results of a TukeyHSD post-hoc test. The peak size of both object types increased with LB concentration.

### S5: Correlation of object count and covered area

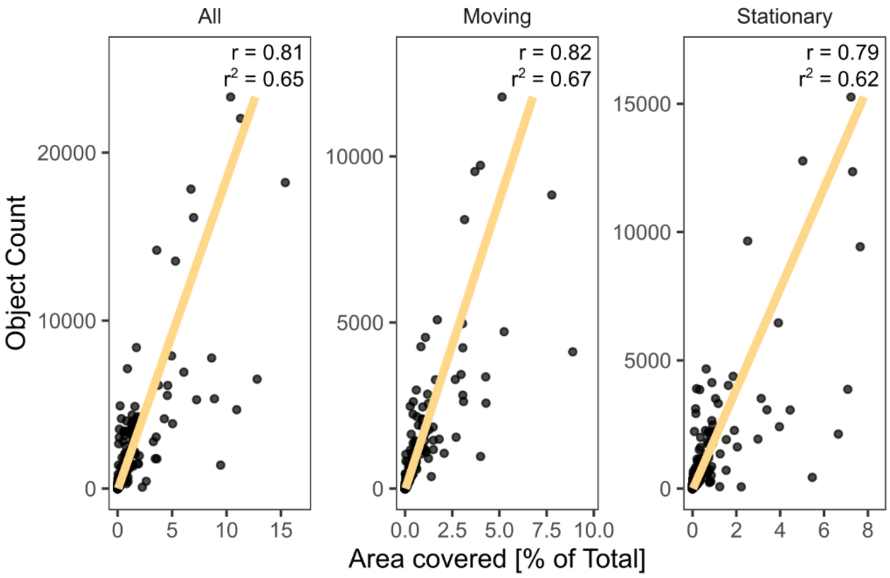

**Figure S4:** Linear regression of covered area and object count of all, moving and stationary objects respectively for all images. Both values are correlated for every object type (r = Pearson correlation coefficient).

**S6: Attached biomass peak size and carrying capacity**

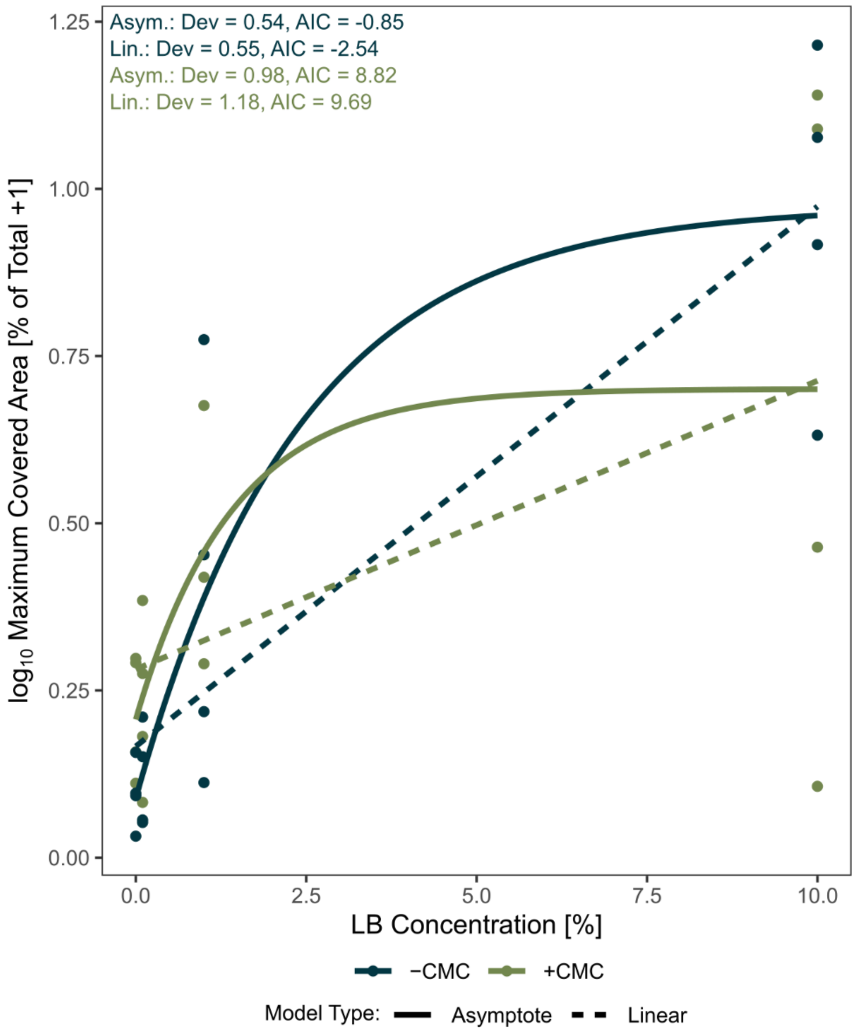

**Figure S5:** Maximum peak size of covered area by stationary objects increased with LB concentration. A linear and a non-linear, asymptotical regression was fit to the data for both CMC groups. Depicted on the plot are the deviance from the data (in absolute values) and the Akaike Information Criterion (AIC) respectively. Both model types show similar fits, with the asymptotical model always having lower deviation and a lower AIC for +CMC. This suggests that the chips were close to the carrying capacity with 10% LB and that adding more would not necessarily lead to a linear increase of the peak size.

**S7: Replicate Comparison Covered Area**

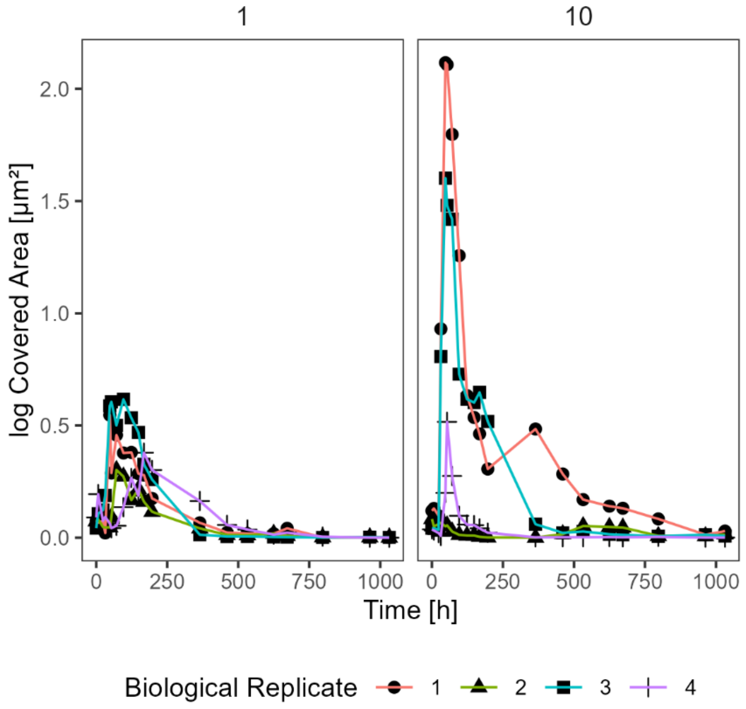

**Figure S6:** Log-transformed covered area of the +CMC chips for the different biological replicates for 1 and 10% LB. Shapes can be directly compared to Figure 7b and c. This reveals a more consistent response across replicates for 1%, while in 10% two replicates (2 and 4) show a lower maximum peak compared to the other two, and these are the same replicates with higher normalized Congo Red fluorescence (Fig. 8b).

**S8: Object size and circularity**

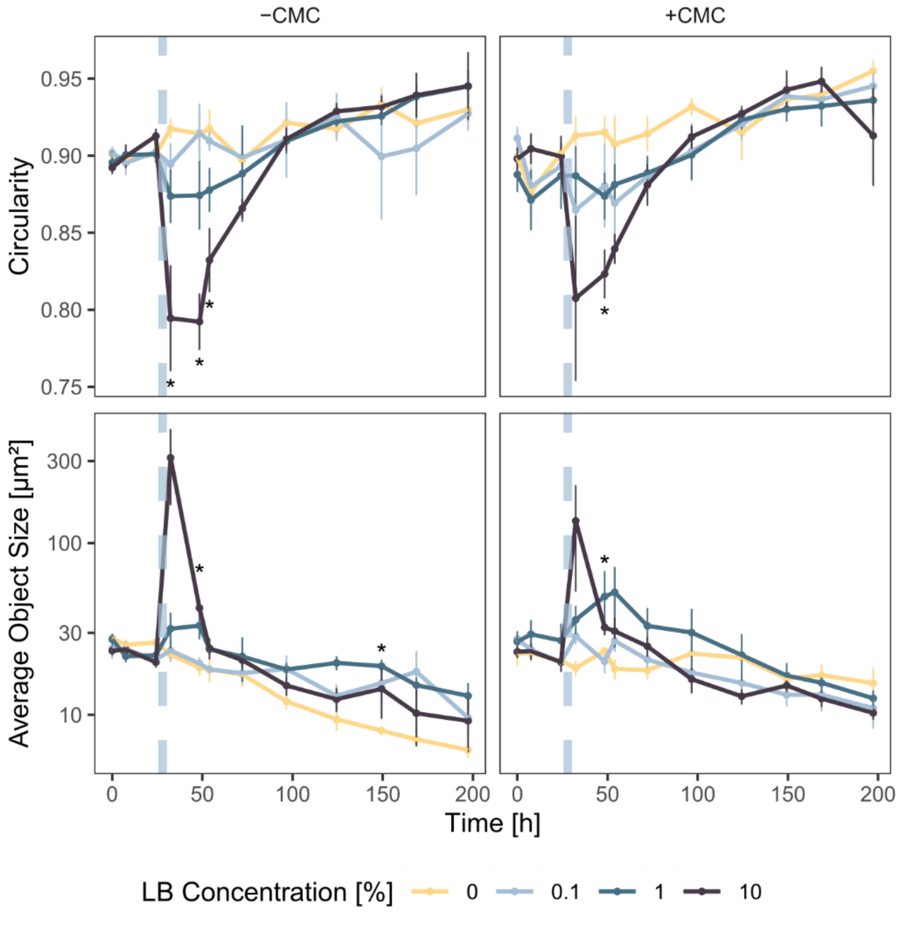

**Figure S7:** Average circularity and object size (mean ± SE) in the first 200 hours of the experiment for the +CMC and -CMC chips and all LB concentrations respectively. The dashed line indicated the priming event. Asterisks represent significant differences of one LB combination compared to the others at the same timepoint, tested with a Welch-test or Mann-Whitney U-test, depending on normality of the data (α = 0.05). The data shows that circularity was minimal while average object size was maximal with 10% LB shortly after the substrate pulse but then reduced to values comparable with the other concentrations again afterwards.

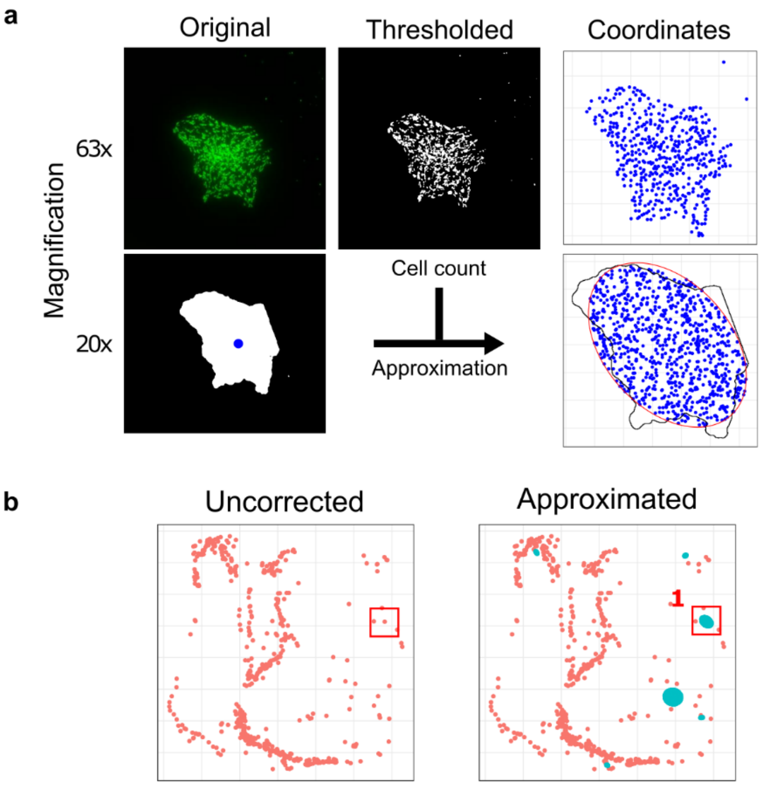

**Figure S8:** Example visualizing the correction employed for large objects that could not be resolved to individual-cell level with the microscopy settings used in the main experiment (a). These objects imaged at 20x were represented by a single coordinate (blue dots), skewing the spatial clustering results. Nine images of two colonies each were taken with higher resolution at multiple timepoints, allowing us to estimate the cell count in these objects through image analysis (upper row of images). We wrote an algorithm in R that takes every large object (with an area larger than one standard deviation from the average of all objects) in the dataset and randomly distributes the expected number of cells in the best-fitting ellipse around the object (lower row of images). This approximates the distribution of bacteria in the 20x images and reduces the bias in the clustering analysis. The planar point pattern of observed objects before and after the correction is shown in b. The red rectangle represents the position of the example object shown in a.

**S9: Spatial Organization**

The spatial organization of objects was compared to complete spatial randomness (CSR) by calculating Ripley's K function (Ripley, 1988) using R package "spatstat" (3.3-1, Baddeley et al., 2015). This approach compares the number of points around a given radius around a specific point and compares it to the number of points that would be expected, given the average density of points. If the derived value is above the expected value, the measured objects exhibit clustering, if below, they are dispersed. Coordinates were converted to planar point patterns with function ppp(). We extracted the shape of the substrate patch as a polygon from Fiji and used that as the window for each point pattern. Using a maximum radius of 50 µm with step size 0.05 and Ripley's correction, the linearized K value for inhomogenous data was calculated for each point pattern with function Linhom() within the envelope() function to obtain confidence intervals. This resulted in a plot similar to that shown in Fig 1j, with the actual L values plotted together with L values that would be expected under CSR. The deviation from CSR was tested statistically with the Diggle-Cressie-Loosmore-Ford test using function dclf.test() with 99 simulations. We quantified the total difference (= area between curves) when the measured value was outside of the CSR upper confidence interval. Negative values (significant dispersion) did not occur. We then summarized this measurement across the entire time-series for each chip to get an idea of overall spatial organization. A caveat of this method is the use of object coordinates, which shrinks large cell clusters down to a single point, with no points in its direct vicinity. To circumvent this bias, we took more detailed images of *B. subtilis* colonies growing in chip 11 with a 63x air objective (HC PL Fluotar 63x/0.90, Leica Microsystems) and calculated the average number of cells per µm in such a colony (see Fig. S8), which came out to be 0.22 ± 0.03 cells * $\mu m^{-2}$. We then wrote an algorithm in R that takes objects larger than one standard deviation above the average object size and randomly places the estimated number of cells at coordinates within the optimal ellipse fit to the object by Fiji, making sure no coordinates were doubled. Clustering was then assessed with this corrected data. No other analysis made use of this simulated data.

**S10: Congo Red Fluorescence**

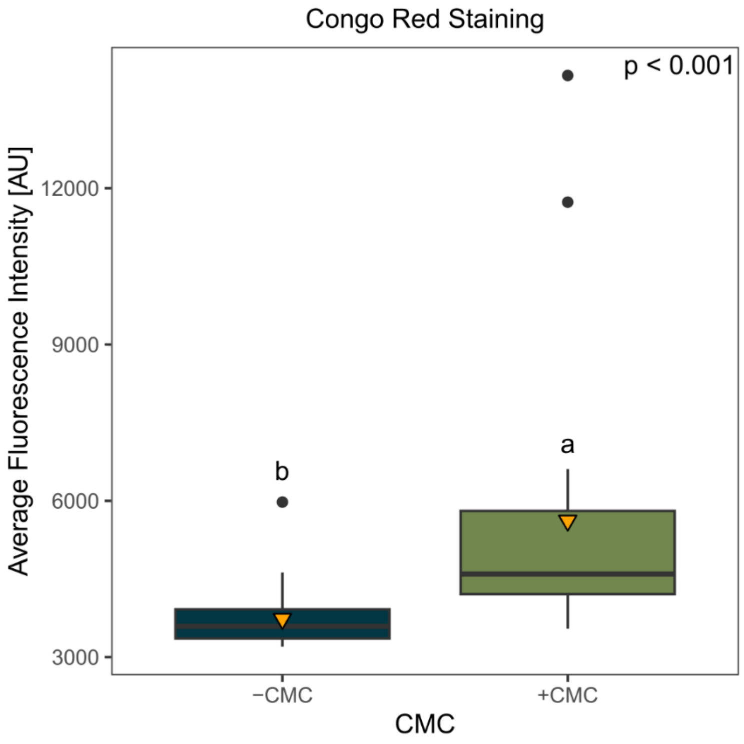

**Figure S9:** Boxplot (median and interquartile range) showing the average Congo Red fluorescence intensity of all images taken at the endpoint. The chips with CMC had significantly higher fluorescence values (Wilcoxon z = -1149.29, d = 0.97), indicating that mostly the remaining CMC was stained instead of a compound of the M9-agarose gel.

**S11: Congo Red fluorescence model**

To explain the differences in Congo Red fluorescence of the CMC-substrate patches, we used linear modelling (R core package “lm”). We first did some tests of different predictor combinations and transformations and compared models by their AIC. Multicollinearity was assessed by calculating variance inflation factors (VIF) using function vif() of package “car” (Fox & Weisberg, 2019). Final initial predictors were LB concentration, average circularity, deviation from CSR and proportion of stationary objects (all averaged over the entire time-series for each chip). As the biomass measurements of covered area, maximum covered area, total object count and average object size were highly correlated (see Table S3), we collapsed them into principal components using function prcomp() and used the first two axis as additional predictors. All predictors and the response were $\log_{10}$-transformed. Predictors were additionally centered and scaled before modelling or collapsing. The full model formula was:

Fluorescence ~ LB concentration + PC1 + PC2 + average circularity + average deviation from CSR + average proportion of stationary objects + average size of objects

Using stepwise elimination of predictors through the step() function, we ended up with a model that best explained the Congo Red fluorescence in our +CMC chips. Normality of model residuals was checked visually and with a Shapiro-Wilk test ($W = 0.96$, $p = 0.73$).

**Table S3: Correlations of biomass-related parameters and their loadings in the final two principal components they were collapsed into**

| Correlations and PCA loadings | Average covered area | Average object count per µm² | Maximum covered area | PC1 | PC2 |
|---|---|---|---|---|---|
| Average covered area | 1 | 0.99 | 0.99 | 0.586 | 0.124 |
| Average object count per µm² | | 1 | 0.99 | 0.576 | 0.632 |
| Maximum covered area | | | 1 | 0.570 | -0.765 |

**S12: CMC decompostition and stain test**

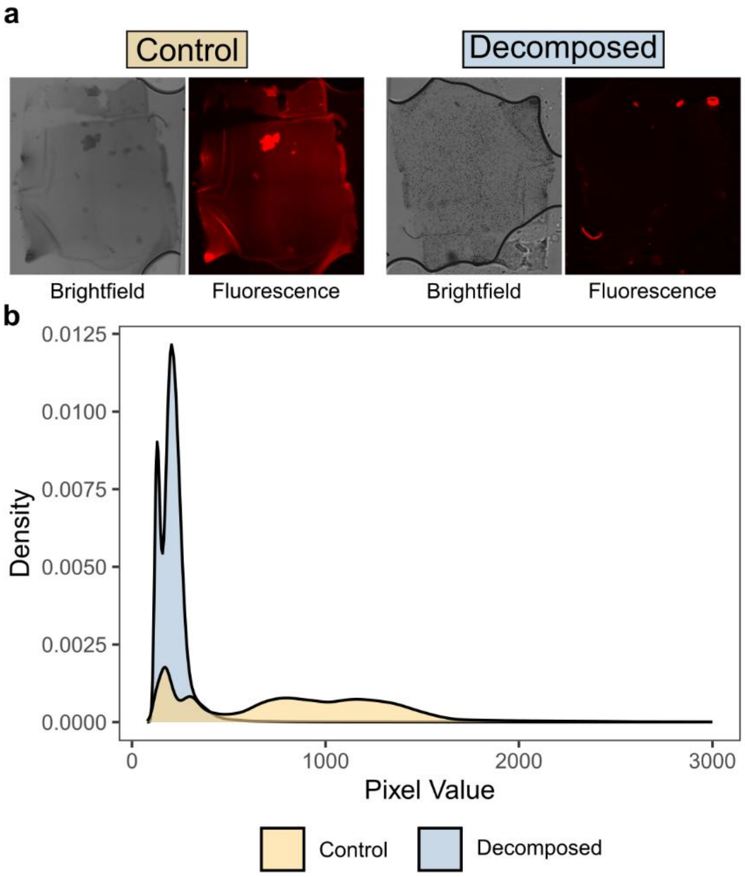

**Figure S10:** Control experiment to verify that CMC decomposition decreases the Congo Red fluorescence of the substrate patch. Two chips were manufactured in the same way as in the main experiment. One was filled with M9 and the other with M9 supplemented with 5 g/l purified cellulase from *Trichoderma viride* (Cellulase Y-C, MP Biomedicals). Chips were stained and imaged (with the same settings as in the main experiment) after 6 hours of incubation at room temperature. Decomposition clearly decreased the overall fluorescence, visible by eye in the fluorescence images (a) and in the histogram of pixel intensities (b), which shows a narrower distribution around lower values in the decomposed gel. Pixel values higher than 3000 were negligible and removed from the plot to increase visibility of the lower regions.

**S13: Carbon approximation**

Nine µl of 2% CMC were added for each gene frame, in the end 1/9$^{th}$ of that ended up in each chip. Assuming a C% of 40% (glucose), we added about 8 µg C to each chip in CMC. Assuming a carbon content of about 5.86 g C * l-1 for LB medium (see Table S6) and an added volume of about 2 µl of each dilution, we can calculate how much C was added to each chip in relation to the CMC-C (Table S5). Carbon in CMC and added medium was much higher than in the initial biomass. We observed about 260 objects in the focal plane after the inoculation. Assuming an average carbon content of a single cell of 12.4 fg (Fukuda et al., 1998), even if the total population in each chip was 10000 cells, total C in the initial biomass would be below 1 ng, so less than 10% of the added C in the lowest LB concentration.

**Table S5:** Proportions of added carbon

| Dilution | µg C * µl -1 | µg C added | % of CMC in chip |
|---|---|---|---|
| 10 | 0.586 | 1.172 | 14.65 |
| 1 | 0.0586 | 0.117 | 1.47 |
| 0.1 | 0.00596 | 0.012 | 0.15 |

**Table S6:** Approximation of carbon content in LB medium

| Compound | | | | | | | C% | g * l$^{-1}$ | g C * l$^{-1}$ |
|---|---|---|---|---|---|---|---|---|---|
| Yeast extract | | | | | | | **36.94**[1] | **5** | **1.847** |
| | Amino Acid (AA) | Formula | g * mol$^{-1}$ | C [%] | g AA in 100g[2] | g C in 100g | | | |
| | Alanine | C3H7NO3 | 89.09 | 40.41 | 2.81 | 1.16 | | **10** | **4.016** |
| | Cysteine | C3H7NO2S | 121.16 | 29.71 | 0.4 | 0.12 | | | |
| | Histidine | C6H9N3O2 | 155.1546 | 46.41 | 2.29 | 1.06 | | | |
| Tryptone | Lysine | C6H14N2O2 | 146.19 | 49.25 | 6.51 | 3.21 | **40.16** | | |
| | Proline | C5H9NO2 | 115.13 | 52.12 | 8.65 | 4.51 | | | |
| | Tryptophan | C11H12N2O2 | 204.23 | 64.63 | 1.05 | 0.68 | | | |
| | Arginine | C6H14N4O2 | 174.2 | 41.33 | 3.31 | 1.37 | | | |

| | | | | | | | | | |
|---|---|---|---|---|---|---|---|---|---|
| | Glutamic acid | C5H9NO4 | 147.13 | 40.78 | 18.7 | 7.63 | | | |
| | Isoleucine | C6H13NO2 | 131.17 | 54.89 | 4.48 | 2.46 | | | |
| | Methionine | C5H11NO2S | 149.21 | 40.21 | 2.35 | 0.94 | | | |
| | Serine | C3H7NO3 | 105.09 | 34.26 | 5.08 | 1.74 | | | |
| | Tyrosine | C9H11NO3 | 181.19 | 59.61 | 1.86 | 1.11 | | | |
| | Aspartic acid | C4H7NO4 | 133.1 | 36.06 | 6.52 | 2.35 | | | |
| | Glycine | C2H5NO2 | 75.07 | 31.97 | 1.79 | 0.57 | | | |
| | Leucine | C6H13NO2 | 131.17 | 54.89 | 7.63 | 4.19 | | | |
| | Phenylalanine | C9H11NO2 | 165.19 | 65.38 | 4.09 | 2.67 | | | |
| | Threonine | C4H9NO3 | 119.1192 | 40.30 | 3.91 | 1.58 | | | |
| | Valine | C5H11NO2 | 117.151 | 51.22 | 5.51 | 2.82 | | | |
| | | | | | | | | **Total** | **5.863** |

[1]Zapata-Vélez & Trujillo-Roldán (2010)

[2]Amino acid composition of Tryptone according to GRiSP Research Solutions, obtained from https://khimexpert.com/wp-content/uploads/2018/12/GCM23-Tryptone.pdf on Dec. 15, 2025.

### S14: Supplementary Methods

#### Bacteria and Culture Conditions

Green fluorescent protein-expressing *B. subtilis* strain 4819 (genotype PS-216-amyE::Pveg(+1/+8)_R0_sfGFP_spec) was kindly provided by Dr. S. Pollak. This is a genetically modified wild type (Durrett et al., 2013) that expresses GFP instead of an alpha-amylase. It differs from most laboratory *B. subtilis* strains because in comparison to those, it did not lose its ability to produce biofilm (Durrett et al., 2013; Omer Bendori et al., 2015). As pre-experiments have shown that this strain only grows in M9 minimal media when tryptophane is present, all M9 used in this experiment (for the casting of agar gels or dilution of LB medium) is supplemented with 20 mg * $l^{-1}$ tryptophane (Sigma-Aldrich). Growth (assessed with optical density at 600 nm, measured with a plate reader (Tecan Spark)) from this concentration of tryptophane alone was negligible compared to when another C source was added (see also the object count and covered area data of the -CMC/0% LB treatment of Figures 2 and 4), so we expect no influence of this addition on our results.

#### Chip fabrication

The chamber design and microfluidic master wafer was kindly provided by Dr. M. Palatzinsky. To obtain the incubation chambers, 8 g of 10:1 Polydimethylsiloxane-mix (PDMS:Curing agent, Sylgard) was poured on top of the cleaned silicium wafer. It was then degassed in an evacuated desiccator for 30 minutes, followed by curing of the mixture at room temperature overnight. The ~2 mm thick chips were then cut out with a scalpel. To help with inoculation, interfacing and reducing evaporation, small PDMS bricks were attached to the top of the in-and outlets by activating both PDMS surfaces for 10 seconds in a plasma oven (Zepto MHz, Diener electronic), followed by baking on a hotplate set to 80°C for 4 minutes. In-and outlets were then punched using a biopsy puncher (WellTech Rapid-Core, 0.75 mm). Finished PDMS chips were stored feature-side up in closed plastic petri dishes until further usage.

Carboxymethylcellulose-agar was produced by slowly stirring 2% w/v CMC (Sigma-Aldrich) in M9 minimal medium overnight until it was completely dissolved, followed by the addition of 1.5% w/v agarose (Roth) and autoclaving. Before casting of the agar layer, the mixture was melted in a microwave and then kept on a hotplate set to 75°C in a laminar flow hood. Gene frames (1 x 1 cm, Thermo Scientific) were attached to glass coverslips (24 x 60 mm, Marienfeld) and shortly heated on the same hotplate as the CMC-agar. Using reverse pipetting, 10 µl of the agar mixture was then pipetted onto the gene frame and evenly spread using the pipette tip. After cooling, a thin and brittle transparent layer of CMC-agar was formed within the gene frame. Using a scalpel, the agar layer was scored carefully in a checkerboard pattern with rectangles of ~ 2 x 4 mm, followed by careful removal of the gene frame and excess agar film using scalpel and tweezers, leaving only the central rectangle attached to the coverslip. The area around this CMC-agar patch was then carefully cleaned using a lint-free paper towel soaked in etOH. For the M9 controls, the same procedure was done using CMC-free M9 agar.

The PDMS chambers were then attached to the glass coverslips, yielding the final decomposer chips. For this, the glass coverslips with the substrate patches were surface-activated (agar side up) in a plasma oven for 60 seconds. Then, the feature side of the PDMS chips was activated for 10 seconds. The activated surfaces were then carefully aligned manually so that the agar patch was right in the middle of the chamber and put in contact. After carefully pressing down on the chips with tweezers to ensure contact without collapsing the features, the finished chips were baked on a hotplate for 4 minutes at 80°C. The assembled chips were then flushed with 70% etOH using a syringe pump (SPLab12, Baoding Shenchen) for 1 h at a flow rate of 1 µl * $min^{-1}$ to sterilize them. Afterwards, they were flushed with sterile M9 minimal medium for 24 h at the same flow rate. This was necessary to ensure sterility and flush out any remaining debris from the fabrication process. The liquid-filled chips were then sealed by covering the in/outlets with small pieces of autoclaved aluminum foil and transparent scotch tape. They were stored horizontally in 50 ml falcon tubes containing a paper tissue wetted with autoclaved MQ (further called "incubation tubes") until the start of the experiment. This was necessary because pre-tests have shown the chips to dry out quickly otherwise.

**Experimental Setup and Incubation**

All work concerning the handling of live bacteria was done in a laminar flow hood. Similarly, inoculated chips were only unsealed in a flow hood. Bacteria were transferred to a fresh plate 24 hours before start of the incubation. Four colonies were transferred into a 2 ml reaction tube each that contained 1 ml of 0.1% LB diluted in M9 minimal media and gently vortexed. LB was added here to ensure establishment of colonies in the chip in the following acclimatization period. Each one of these tubes was further treated as a biological replicate. Each suspension was diluted to an $OD_{600}$ of 0.001 to ensure similar initial cell densities.

Washed chips were UV-irradiated in a laminar flow hood for 1 hour. Remaining liquid was flushed out by forcing sterile laminar flow hood air through the chamber using a 10 µl pipette. Chambers were then carefully filled with cell suspension using a 10 µl pipette, making sure not to introduce any large air bubbles in the system. After inoculation, chips were sealed again and stored horizontally in the incubation tubes at 30 °C. For each biological replicate, 4 chips with CMC-agar and 4 chips with M9-agar were inoculated this way (32 chips in total). Four additional chips each were just inoculated with M9 minimal medium and kept in the incubator as sterile controls to assess CMC decomposition in the end (see below).

After an acclimatization period of 48 hours, non-control chips were given a priming pulse of different concentrations. One chip of each biological replicate of each CMC treatment was flushed with 0%, 0.1%, 1% and 10% of LB medium diluted in M9 minimal medium. The pulse was delivered using sterile syringes (Braun Omnifix 5 ml), a syringe pump (SPLab12, Baoding Shenchen) and ethanol-sterilized tubing (PE 50/10, Warner Instruments) in a laminar flow hood. The flow rate was set to 0.7 µl * $min^{-1}$, which was the lowest possible setting that delivered enough force to push liquid through the chamber while minimizing the direct impact on the bacteria. The flow was maintained for 5 minutes, ensuring that the chamber volume (about 1.5 µl) was replaced once by the solution. Chips were then sealed and incubated in their tubes again and were ready for long-term imaging.

**Image Aquisition**

Chips were taken out of their incubation chambers and imaged at multiple timepoints. Table S7 shows an overview of the timeline and intervals between timepoints. Images were acquired in 16 bit using an inverted wide-field fluorescence microscope (Leica Thunder Imager) equipped with a Leica K8 camera (Leica Microsystems) and an incubation chamber (Okolab Microsystems). The image acquisition workflow was the same for each chip and timepoint during the incubation.

**Table S7: Overview of timeline and sampling intervals**

| Timepoints | Hours since start | Intervals |
|---|---|---|
| T1-T8 | 0-96 | Twice daily, 8h apart |
| T9-T12 | 96-200 | Every 24h |
| T12-T20 | 200-1000 | Irregular, every 100-200h |

Each chip was separately taken out of the incubation chamber and put onto the microscopy stage, which was set to the same temperature as the long-term incubator (30 °C). Acquisition areas were defined for each chip and kept constant across all timepoints. They covered the whole agar patch area plus some of the chamber edge to help with alignment later. Areas were roughly aligned with pre-determined static points on the edge of the chamber, to ensure that they were at similar positions relative to the chip for every acquisition. The focal plane was set so that the first cells that are visible are in focus, since those are the cells on the surface of the agar films as the microscope was inverted. The LEICA application suite software (LAS X, version 3.9.1.28433) allows the definition of multiple focus points in an area. This was done to interpolate the focal plane between different regions on the agar film, which might be otherwise out of focus as the agar surface is not necessarily completely flat. A tilescan of the acquisition area was then performed using a 20x air objective (HC PL Fluotar 20x/0.55, Leica Microsystems) and automatically stitched together by the software. GFP in the cells was then imaged with a base light intensity of 100% and 200 ms exposure time (475/535 nm excitation/emission). Intensity and acquisition time had to be adjusted when a lot of cells were present to avoid overexposure. The used image acquisition parameters were recorded for every image and used for thresholding (see below). Two images with the same settings were taken 30 seconds apart. After imaging, the chip was returned to its incubation tube and placed back in the long-term incubator. From time to time a few drops of autoclaved MQ were added to the paper towel to ensure sufficient moisture in the falcon tube.

**Image Processing**

Image analysis was done using Fiji (2.16.0/1.54p) with custom scripts to semi-automate some processes. In a first step, the fluorescence images were converted to 8 bit. For each chip, each image in a time-series was aligned to the first image (taken at T1) using the line ROI alignment tool. For this, the images were duplicated and had their contrast increased, making the chamber edge easily visible. Then, a straight vertical line was drawn across the edge in both the T1 and Tx image, x being any other timepoint (2-20). The line was then transferred to the original image. The tool then overlays these lines and shifts the coordinates of the Tx image to match the one of T1. The same was then done using a horizontal line on the upper or lower edge of the visible agar patch. A polynomial region of interest was then drawn on the agar patch area with a distance to the edge to exclude edge effects, copied to each aligned image and cropped. As the agarose substrate was subject to swelling and shrinking over the course of the experiment, the area had to be slightly adjusted for each timepoint. In some cases, the agarose substrate was folded over itself in some areas during the fabrication process, resulting in focal planes with a too large difference for the focus point interpolation. We therefore drew the polygon only around the area with the in-focus cells. Measurements were normalized to the polygon area to compensate for this difference.

Next, images were thresholded. We used the same threshold on images acquired with the same settings (exposure time + light intensity combination). Using the Otsu thresholding algorithm, each image was first processed using the automatically detected threshold. In cases where there were no bacteria present, the automatic detection picked up on background noise, resulting in a low threshold and producing false positives. By visual inspection of the histogram of thresholds, a minimum threshold was therefore set for each combination of exposure time and light intensity. We then calculated the mean threshold of all automatically set thresholds above the minimum threshold for each image acquisition-settings group to finally process each image. This method improved comparability between images while minimizing bias introduced by manual threshold setting.

In a small number of cases, auto-fluorescent dust on top of the PDMS chamber resulted in artifacts in the images. These were identifiable because they did not change their position or shape over the course of the experiment. These artifacts were manually removed from the images by setting their pixel values to 0, taking care not to delete any actual cells.

For our research question, we were interested in the sessile cells on top of the agar patch. However, as *B. subtilis* is flagellated, we observed a lot of moving cells as well. To separate moving from stationary cells, we employed the TrackMate (Ershov et al., 2022) plugin of FIJI. We were able split each image (consisting of two images 30 seconds apart, see above) into an image containing objects that stayed in the same position within 30 seconds (= stationary cells for our purpose) and one with those that either moved or vanished (= moving cells). After some testing, the plugin was configured with the mask detection algorithm (as the images were already thresholded) and a maximum linking and gap-closing distance of 200 µm. For dense images with thousands of objects, the distance was reduced to 50 µm to reduce computation time. Very large stationary objects had to be excluded in some images as they produced false positives. The “simple Linear Assignment Problem” (LAP) algorithm was chosen to detect moving objects. Once tracking was done, all objects that moved more than 5 µm, which was the approximate average Feret diameter of objects in a test set of images, were deemed moving and exported into a new image. This threshold was necessary to remove false positives introduced by minor shifts between the two images. By simply subtracting the newly created image containing the moving objects from the original image containing all objects, we were able to produce an image of stationary objects. Comparing the sum of extracted moving and stationary objects to the sum of total objects detected in the original image containing both, we calculated a mean loss of 0.051 ± 0.18% of objects, which we see as an acceptable error rate for this approach.

Total object count, covered area and mean circularity were finally extracted from each aligned, thresholded and tracked image for further data processing. Individual object coordinates, size and circularity was additionally exported.

**Cellulose Decomposition Assessment**

After the final timepoint, chips were flushed with 5 µl of phosphate-buffered saline (PBS) and then filled with ice-cold absolute ethanol (Supelco) in a laminar flow hood. They were then sealed and stored at 4°C until further processing. Congo Red staining was performed using the protocol of Romano et al. (2013), modified to our purpose. A 0.5% w/v CR (Sigma-Aldrich) solution was prepared in MQ. Each chamber was then filled with the CR solution and incubated at room temperature for 15 minutes. Then, the stain was deactivated using 1M NaCl solution, followed by 3-4 flushes with MQ until no dye was visible in the outflow. The stained gels were then observed under the microscope using the same setup as above (200 ms acquisition time, 100% intensity, 575/590 nm excitation/emission). The images of Congo Red stained substrate patches were cropped to the substrate area, excluding oversaturated areas with clear artifacts. Images were then down-sampled by 50% to reduce file size using the “scale” command in Fiji. A list of the intensity values of these down-sampled pixels was then exported. This was done instead of just exporting the average intensity because we wanted to investigate co-location of bacteria and decomposed areas, however this yielded no significant results.

**Calculations and Statistics**

Statistical tests were performed with the respective function of R package “rstatix” (0.7.2, Kassambara, 2023) if not mentioned otherwise. The effects of the manipulated factors CMC and LB concentration on non time-resolved variables were assessed using ANOVA (function aov() of core R package ”stats”) or Kruskal-Wallis tests if ANOVA prerequisites homogeneity of variance and/or normality of residuals were violated (checked with Levene’s and Shapiro-Wilk tests respectively). If assumptions of a two-way ANOVA were not met, both factors were combined into a dummy variable and used in an overall Kruskal-Wallis test. Differences between groups were assessed with the post-hoc tests Tukey honest significant differences or Dunn ‘s test with Benjamini-Hochberg correction respectively. If applicable, a two sided Welch-T test for normally distributed data or a Mann-Whitney-U test for non-normal data was used to test differences between two groups (for example, Fig. 7a). For all tests, effect sizes were calculated and reported for significant effects (Cohen’s D for Welch/MWU, $\eta^2$ for ANOVA/Kruskal-Wallis).

### References

Arel-Bundock, V., Greifer, N., & Heiss, A. (2024). How to Interpret Statistical Models Using marginaleffects for *R* and *Python*. *Journal of Statistical Software*, *111*(9). https://doi.org/10.18637/jss.v111.i09

Baddeley, A., Rubak, E., & Turner, R. (2015). *Spatial Point Patterns: Methodology and Applications with R*. Chapman and Hall/CRC.

Durrett, R., Miras, M., Mirouze, N., Narechania, A., Mandic-Mulec, I., & Dubnau, D. (2013). Genome Sequence of the Bacillus subtilis Biofilm-Forming Transformable Strain PS216. *Genome Announcements*, *1*(3), e00288-13. https://doi.org/10.1128/genomeA.00288-13

Ershov, D., Phan, M.-S., Pylvänäinen, J. W., Rigaud, S. U., Le Blanc, L., Charles-Orszag, A., Conway, J. R. W., Laine, R. F., Roy, N. H., Bonazzi, D., Duménil, G., Jacquemet, G., & Tinevez, J.-Y. (2022). TrackMate 7: Integrating state-of-the-art segmentation algorithms into tracking pipelines. *Nature Methods*, *19*(7), 829–832. https://doi.org/10.1038/s41592-022-01507-1

Fox, J., & Weisberg, S. (2019). *An R Companion to Applied Regression* (Third). Sage. https://www.john-fox.ca/Companion/

Fukuda, R., Ogawa, H., Nagata, T., & Koike, I. (1998). Direct Determination of Carbon and Nitrogen Contents of Natural Bacterial Assemblages in Marine Environments. *APPLIED AND ENVIRONMENTAL MICROBIOLOGY*, *64*(9), 3352–3358. https://doi.org/10.2331/suisan.77.134

Kassambara, A. (2023). *rstatix: Pipe-Friendly Framework for Basic Statistical Tests*. https://rpkgs.datanovia.com/rstatix/

Omer Bendori, S., Pollak, S., Hizi, D., & Eldar, A. (2015). The RapP-PhrP Quorum-Sensing System of Bacillus subtilis Strain NCIB3610 Affects Biofilm Formation through Multiple Targets, Due to an Atypical Signal-Insensitive Allele of RapP. *Journal of Bacteriology*, *197*(3), 592–602. https://doi.org/10.1128/JB.02382-14

Ripley, B. D. (1988). *Statistical Inference for Spatial Processes*. Cambridge University Press.

Romano, N., Gioffré, A., Sede, S. M., Campos, E., Cataldi, A., & Talia, P. (2013). Characterization of Cellulolytic Activities of Environmental Bacterial Consortia from an Argentinian Native Forest. *Current Microbiology*, *67*(2), 138–147. https://doi.org/10.1007/s00284-013-0345-2

Simpson, G. L. (2024). gratia: An R package for exploring generalized additive models. *Journal of Open Source Software*, *9*(104), 6962. https://doi.org/10.21105/joss.06962

van Rij, J., Wieling, M., Baayen, R., & van Rijn, H. (2022). *itsadug: Interpreting Time Series and Autocorrelated Data Using GAMMs.* (Version 2.4.1) [Computer software].

Wood, S. (2017). *Generalized Additive Models: An Introduction with R* (2nd ed.). Chapman and Hall/CRC.

Zapata-Vélez, A. M., & Trujillo-Roldán, M. A. (2010). The lack of a nitrogen source and/or the C/N ratio affects the molecular weight of alginate and its productivity in submerged cultures of Azotobacter vinelandii. *Annals of Microbiology*, *60*(4), 661–668. https://doi.org/10.1007/s13213-010-0111-7
